# Spontaneous mutations and transmission distortions of genic copy number variants shape the standing genetic variation in *Picea glauca*

**DOI:** 10.1101/103796

**Authors:** Atef Sahli, Isabelle Giguére, Jean Bousquet, John MacKay

## Abstract

Copy number variations (CNVs) are large genetic variations detected among the individuals of every multicellular organism examined so far. These variations are believed to play an important role in the evolution and adaptation of species. In plants, little is known about the characteristics of CNVs, particularly regarding the rates at which they are generated and the mechanics of their transmission from a generation to the next. Here, we used SNP-array raw intensity data for 55 two-generations families (3663 individuals) to scan the gene space of the conifer tree *Picea glauca* (Moench) Voss for CNVs. We were particularly interested in the abundance, inheritance, spontaneous mutation rate spectrum and the evolutionary consequences they may have on the standing genetic variation of white spruce. Our findings show that CNVs affect a small proportion of the gene space and are predominantly copy number losses. CNVs were either inherited or generated through *de novo* events. *De novo* CNVs present high rates of spontaneous mutations that vary for different genes and alleles and are correlated with gene expression levels. Most of the inherited CNVs (70%) are transmitted from the parents in violation of Mendelian expectations. These transmission distortions can cause considerable frequency changes between generations and be dependent on whether the heterozygote parents contribute as male or female. Transmission distortions were also influenced by the partner genotype and the parents’ genetic background. This study provides new insights into the effects of different evolutionary forces on copy number variations based on the analysis of a perennial plant.

## Introduction

Copy number variations (CNVs) refer to specific sequences that are present in variable numbers in the genome of different individuals. CNVs are among the least studied genetic variations despite their abundance in natural populations (Blackburn et al. 2013; Chain et al. 2014; Jakobsson et al. 2008; Kato et al. 2010; Mills et al. 2011; Tan et al. 2012), the considerable proportion of the genome they affect (12-15% of the human genome; Redon et al. 2006; Sebat et al. 2004) and their impacts on gene function (Conrad et al. 2010; Debolt 2010; Korbel et al. 2009; Mills et al. 2011), gene expression (Schlattl et al. 2011; Stranger et al. 2007) and downstream phenotypes (Beckmann, Estivill, and Antonarakis 2007; McCarroll and Altshuler 2007). Genome-scale identifications of CNVs have been reported in many species including some plants thanks to the rapid development of genotyping and sequencing technologies in the last two decades (Saxena et al. 2014; Zemienko et al. 2014); however, much less is known about their generation and transmission through generations and the impact of evolutionary forces upon them.

A better understanding of the dynamics of spontaneous mutations will help to elucidate the role of different genetic variations in shaping of genome architecture and evolution, and in adaptation. But the current knowledge is limited to a few species, particularly for CNVs, and estimates obtained to date are often biased due the sampling or method of analysis (reviewed in Katju and Bergthorsson 2013; Itsara et al. 2010). CNVs can be generated by different molecular mechanisms (Chen et al. 2010), are associated with different local features of the genome architecture (Baer, Miyamoto, and Denver 2007), have a large impact on gene structure and function and on fitness (Tang and Amon 2013) and, are frequently induced under stress conditions (Debolt 2010). All of these factors can influence the mutation rates of CNVs across the genome. The rates at which new variants appear in a population is transient and will reach an equilibrium resulting from the action of different evolutionary forces. While purifying selection drives the mutation rate (μ) to lower levels in order to minimize the deleterious effect of disruptive mutations, adaptive selection, drift and the need to reduce the fidelity cost drive μ to higher levels.

Variants generated through *de novo* mutations or introduced through sexual reproduction are expected to be transmitted randomly to the next generation according to Mendelian expectations. Transmission distortions (TD) occur when an allele is preferentially transmitted to the next generation at the expense of alternative alleles. This departure from the Mendelian expectations is observable in the offspring of heterozygote individuals and is often the consequence of disruptive mechanisms operating during the gametic or zygotic stages of development. TDs involving CNVs at responders and/or distorters loci were identified in a few model organisms including the loci om and WSB in mice (Pardo-Manuel De Villena et al. 2000; Didion et al. 2015) and the peel-1 locus in *C. elegans* (Seidel et al. 2011). A more comprehensive analysis of the transmission of CNVs and the occurrence of copy number transmission distortions at the genome scale will help to better understand how genetic variations are transmitted and maintained in a population.

CNVs and their role in evolution and adaptation is poorly understood in non-model organisms. In trees, CNVs have been investigated only in a few species and for a limited number of loci or individuals (Šķipars, Krivmane, and Ruņģis 2011; Neves et al. 2013; Pinosio et al. 2016; Prunier, Caron, and MacKay 2017). In this work, we screened the gene space of white spruce (**Picea glauca**), a perennial outcrossing monoecious gymnosperm with a diploid giga-genome (20 Gb) enriched in repeated sequences (Birol et al. 2013; De La Torre et al. 2014; Warren et al. 2015), to identify genic CNVs and characterize their inheritance and rates of spontaneous generation. The use of mutation accumulation lines is not feasible for most species (especially those with a long generation time) and does not inform on the inheritance of genetic variations; therefore, we used a family based approach instead. We took advantage of the availability of raw intensity data from a SNP-array targeting more than 14000 gene coding sequences that was used to genotype 3663 individuals in 55 two-generation pedigrees (Pavy et al. 2013; Beaulieu et al. 2014). These data allowed us to detect genic CNVs in the genome of *P. glauca* and ask the following questions: 1) Are CNVs abundant and what class of CNVs is prevalent in the gene space? 2) What is the rate of CNV generation and how variable is it between genes and alleles? 3) Is there a link between expression of a gene and its mutation rate? 4) Are CNVs inherited according to Mendelian expectations or are they subject to transmission distortions? 5) What types of evolutionary forces act on CNVs and what are their evolutionary and adaptive consequences on the species?

## Materials and Methods

### Data sets

For the purpose of identifying genic CNVs using a cross-samples approach (Marioni et al. 2007), we selected two subsets of raw intensity data previously generated for SNP genotyping analyses of 55 white spruce families (Pavy et al. 2013; Beaulieu et al. 2014). In the two data sets designated 54F and 1LF, *Picea glauca* trees from two generations pedigrees were genotyped using PGLM3 SNP-array (14140 probes targeting 14058 genes). The design of PGLM3 Infinium SNP-array (Illumina, San Diego, California) and the genotyping protocol are described in (Pavy et al. 2013).

The data set 54F consist of the genotyping data of 54 full-sib families (family size range: 28 to 32) with their respective parents (total of 1650 offsprings + 37 parents). Each of the 37 parents was involved in one to five crosses and was used as male and female indistinctively in different crosses. This data set includes also two technical replicates for 24 trees, genotyped on different arrays for quality control. The data set 1LF correspond to the genotyping data of a single large family. The 1974 offsprings of the ♀77111 × ♂2388 cross were genotyped along with five technical replicates of six individuals and the parents 77111 and 2388 were genotyped 12 times each on separate arrays.

### Copy number inference

X/Y intensities (corresponding to A/B alleles’ probes, respectively) were normalized using Illumina proprietary software Genome Studio V2011.1 (Illumina, San Diego, California). The normalized signal intensity data were then exported for copy number inference using the algorithms PlatinumCNV (Kumasaka et al. 2011) and GStream (Alonso et al. 2013). Both algorithms were used with default parameters except for the call rate (in PlatinumCNV) were a conservative threshold of 0.999 was used instead of 0.99 (default value).

A two-steps approach was applied for quality control. First, CNV calls displaying reproducibility between technical replicates below 95% were excluded. Second, the copy numbers inferred for each individual at each locus by both algorithms were compared. CNV calls showing a consistency between the two algorithms below 95% were excluded. For the two data sets analyzed, on average 60% of the initially called CNVs were retained for further analysis.

### CNVs validation with real time qPCR

Primers for 15 CNV genes and 6 candidate reference genes were designed using Primer3 algorithm (Koressaar and Remm 2007; Untergasser et al. 2012). Primers properties (self-hybridization, hairpin loop formation and dimers formation) were assessed *in silico* using OligoCalc (Kibbe 2007). Hybridization temperature (Tm) and DNA concentration were optimized for each qPCR assay. The high resolution melting curves were inspected for each assay and the amplicons were sequenced to check that each reaction amplify the intended target sequence in the genome.

Six candidate reference genes were selected based on their stable expression in *Picea glauca* (Beaulieu et al. 2013). The software geNorm (Vandesompele et al. 2002) was used to test the copy number stability of the candidate reference genes in our DNA samples, and the gene BT112014 (coding for the eukaryotic translation initiation factor 4e) was selected as reference gene for the validation of CNV calls using qPCR.

To validate the discovered CNVs, quantitative real time PCR was performed for 15 genes with 22 to 44 samples for each gene. qPCR reactions were prepared in 384 micro-well plates using epMotion 5075 automated liquid handler (Eppendorf, Hamburg, Germany). Each 15 μl reaction contained 20 ng genomic DNA, 7.5 μl Qiagen Fast Protocol Master Mix (1x) (Qiagen, Hilden, Germany) and forward/reverse primers (300 nM final concentration). qPCR reactions were run in four replicates on a LightCycler480 instrument (Roche Life Science, Penzberg, Germany) and thermal cycling conditions were 95°C for 5 min followed by 50 cycles of 94°C for 10 s and 62°C for 1 min. At the end of the 50th cycle, a high resolution melting curve was generated as follow: 95°C for 1 min, 40°C for 1 min, and a final step of continuous temperature increase from 55°C to 95°C with a 0.02°C/s ramp rate.

The efficiency of each qPCR reaction was estimated using the linear regression method described in (Boyle, Dallaire, and MacKay 2009). The Cp (Crossing point) values and efficiency estimates were imported in the software REST 2009 (Qiagen, Hilden, Germany) (Pfaffl, Horgan, and Dempfle 2002) for further analysis. Copy number ratios were calculated using the BT112014 gene as reference, the parent of each sample as calibrator and an efficiency correction for each reaction. A randomization test (with 10,000 iterations and a significance level α = 0.05) was used to identify significant differences in copy numbers between samples.

### Pedigree reconstruction

Forty-three pedigrees were selected representing the 54 full-sib families genotyped in the 54F data set. Thirty-three of these pedigrees encompassed half-sib families sharing a common parent (two to five families per pedigree). The remaining 10 pedigrees involved two to four unrelated families each. Pedigree reconstructions, based on Allele Specific Copy Numbers (ASCN) from 23 and 79 loci, were performed separately using the maximum-likelihood method implemented in the software Colony 2.0 (Wang and Santure 2009; Jones and Wang 2010; Wang 2013). The reconstruction of each pedigree was performed in three independent runs (with different seeds to start each run) in order to check the convergence of runs toward the same optimal solution. Medium length runs with allele frequency update and no sibship prior were used. The optimal pedigree configuration was identified using the full-likelihood method. The accuracy of pedigree reconstruction using ASCN genotypes was estimated through the parameters P(FS|FS), P(HS|HS), P(UR|UR), P(PO|PO) defined in Wang and Santure (2009).

## Statistical analyses

Transmission ratio of the copy number allele A0 was estimated as follow:

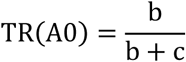

Where b and c are the number of A0 (zero-copy allele) and A1 (one-copy allele) transmissions from a heterozygous parent to his offsprings, respectively. Significant transmission distortions (defined as TR(A0) departures from Mendelian expectations of 0.5) were identified using a two-tailed exact binomial test (α = 0.05). The TR(A0) 95% confidence interval and the test *p-value* were calculated using the function *binom.test* in R (R Core Team 2016).

The distribution of TR(A0) for different families and the distribution of mutation rates (μ_ls_) for different genes were characterized using two approaches i) computation of the Gaussian Kernel Density (GKD) using the function *density* in R (R Core Team 2016) and ii) fitting of a Gaussian Mixture Model (GMM) using the package *mclust* in R (R Core Team 2016). Since both approaches provided similar results, we chose to present only the GKD distributions in this paper.

To examine the potential relationship between the transmission ratios TR(A0) and the genetic distance between parents, we selected 8452 high quality SNPs (Gene Train Score = 0.5 and Call Rate = 1) from the Illumina SNP-array PGLM3 (Pavy et al. 2013). Genotypes for these SNPs are available (Beaulieu et al. 2014) for the 37 parents analyzed in this study. We calculated the genetic distance gd between a pair of parents as the proportion of loci at which the two genotypes being compared were different (Nei and Kumar 2000). The two-dimensional clustering analysis between TR(A0) and gd was conducted using the package mclust in R (R Core Team 2016).

The dependence of Δp (evolution of copy number allele A0 frequency between the parental generation n and the next generation n+1) on TR(A0) and pq (product of the frequencies of copy number alleles A0 et A1 in the parental generation, respectively) values was demonstrated through i) fitting of the data to the model proposed by Chevin and Hospital (2006) and ii) ANOVA using the function aov in R (R Core Team 2016).

The function *cor.test* in R (R Core Team 2016) was used to calculate the one-tailed Pearson correlation coefficient (Cor) between two variables, and associated *p-values*.

## Data availability

SNP-array raw intensity data (X/Y) for the data sets 54F and 1LF, copy number genotypes for CNV genes and qPCR crossing point (Cp) and reaction efficiency (E) data will be submitted to the public data repository Dryad.

## Results

We detected copy number variations (CNVs) in the *P. glauca* gene space by using SNP-array raw intensity data for 14,058 genes in 3663 individuals (for details on the CNV calls, see methods). We characterized the inheritance and estimated the spontaneous mutation rates of CNVs by analyzing 55 two-generation families (54 small families in the 54F data set and one large family in the 1LF data set). Hereafter, we consider a copy number (CN) of two as the normal state for a gene (P. glauca being a diploid organism) and variants (also called non-two-copy genotypes) as homozygous deletions (CN = 0), heterozygous deletions (CN = 1) or copy number gains (CN = 3 or 4).

### Detection and validation of genic CNVs in pedigree populations

We identified CNVs affecting 143 different genes among individuals (Table 1). The genic CNVs detected in each data set represent a small proportion of the 14,058 genes inspected (0.5% on average). Most of the variants (90%) are CN losses (homozygous and/or heterozygous deletions) and only 10% are CN gains (Table 1). No two-way (tri-allelic) CNVs were detected.

**Table 1:**
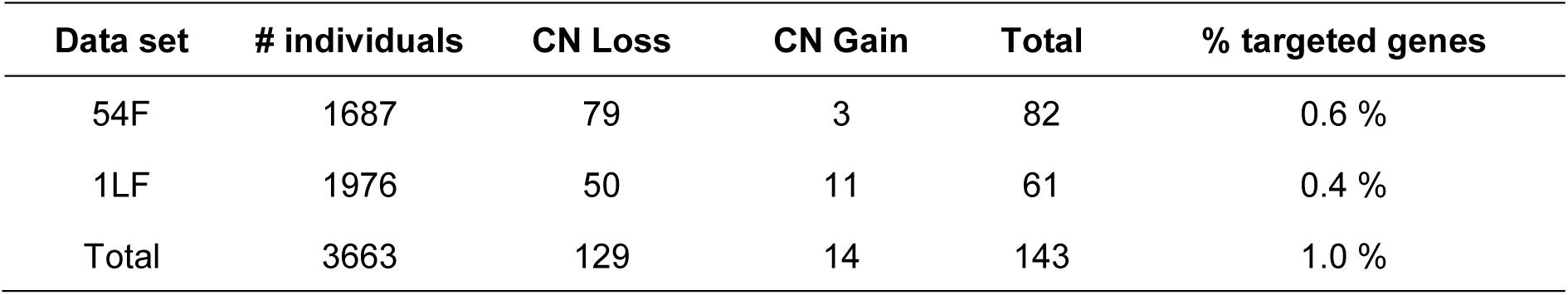
Detected CNVs.

| Data set | # individuals | CN Loss | CN Gain | Total | % targeted genes |
| --- | --- | --- | --- | --- | --- |
| 54F | 1687 | 79 | 3 | 82 | 0.6 % |
| 1LF | 1976 | 50 | 11 | 61 | 0.4 % |
| Total | 3663 | 129 | 14 | 143 | 1.0 % |

We validated the CNV calls using quantitative real time PCR as independent technique. Fourteen out of the 15 tested genes displayed CNVs with both techniques and the estimated False Discovery Rate (FDR) was 6.6%. Careful examination of the gene BT102213 displaying discrepancies between the two technologies showed that while the qPCR primers target a conserved region of the gene, the array probe is located in an LRR1 domain that can be present in one or two copies in different variants of the gene. We were unable to design a probe and primers that target the same region of the gene because of the different technical requirements of the genotyping array and the qPCR. The genotyping accuracy assessed through qPCR was 80% on average but depended on the CNV class. It was high for deletions, i.e. 85% and 87% for heterozygous and homozygous deletions, respectively, and it was low for CN gains (63%) mainly due to a lack of sensitivity of the SNP-array technology for gains (the median sensitivity for CN gain is 20%).

### Classification of CNVs as inherited or *de novo*

CNVs were classified into two categories i) inherited CNVs and ii) *de novo* CNVs to identify the source of the CN variants observed in the offspring generation (Figure 1-A). Inherited CNVs are observed when a non-two-copy genotype is detected in at least one of the parents and among the offspring. *De novo* CNVs on the other hand, are observed when both parents have a two-copy genotype and a non-two-copy genotype is detected among the offspring, which is presumed to result from a spontaneous mutation (loss or gain of a copy). For the 54F data set, 23 (28%) of the identified CNVs were transmitted from parents to their offspring and 59 (72%) of the CNVs were detected as *de novo* events. Each of the families displayed eight inherited CNVs and 23 *de novo* CNVs on average. The remaining genes (51) were maintained at two copies for all the family members (Figure 1-B). The narrow whisker-boxes in figure 1-B show consistent proportions of inherited versus *de novo* CNVs within the different families. A similar profile was observed in the 1LF data set where seven inherited CNVs and 54 *de novo* CNVs were detected (Figure 1-C). This observation indicated that the estimates obtained from small families were not considerably biased. The larger number of *de novo* CNVs identified in the large family is to be expected due to the very large sample size.

**Figure 1:**
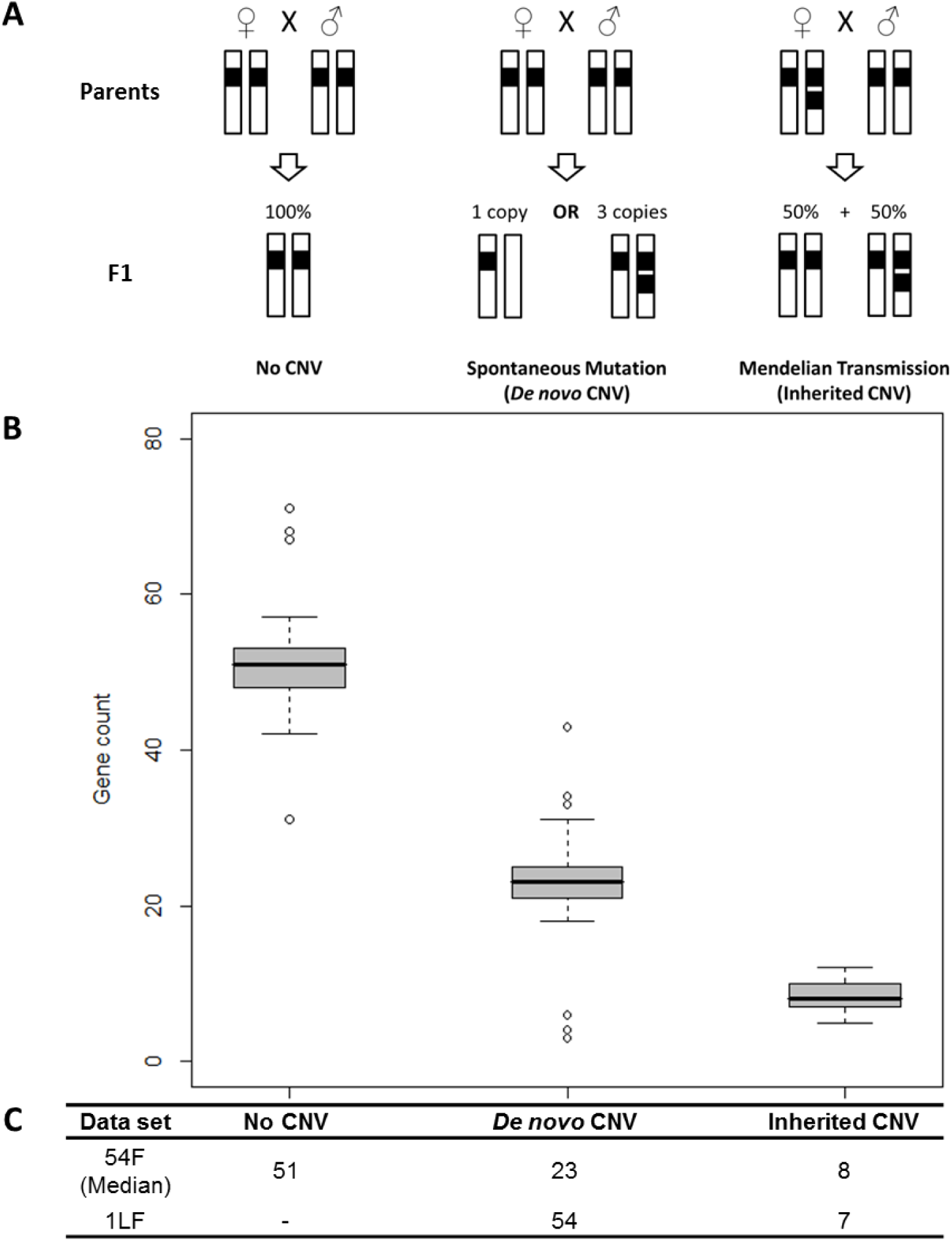
CNVs Classification. Definition of CNV categories according to their status in the parental and F1 generations (A). CNV genes distribution within the three CNV categories for 54 full-sib families (B). Number of *de novo* and inherited CNVs per family (C).

We proceeded to the reconstruction of two-generations pedigrees from the 54F data set using a maximum likelihood approach based on allele specific copy numbers (ASCN). The pedigrees reconstruction using the 23 inherited CNV genes only was achieved with an average accuracy ranging from 91.7 to 95.5% depending on the nature of the relation between individuals (Figure 2). The proportion of dyads correctly inferred was 91.7, 95.5, 92 and 94% for full-sib, half-sib, unrelated individuals and parent-offspring dyads, respectively. In an independent simulation, we used both inherited and *de novo* CNVs genotypes for the reconstruction of the same pedigrees. As might be expected, this decreased the accuracy of full-sib, half-sib and parent-offspring dyads inference and increased the inference accuracy for dyads of unrelated individuals (Figure 2). This result highlights the quality of CNV genotyping using raw data from SNP-arrays and the proper classification of the observed CNVs into inherited and *de novo* CNVs that ensued.

**Figure 2:**
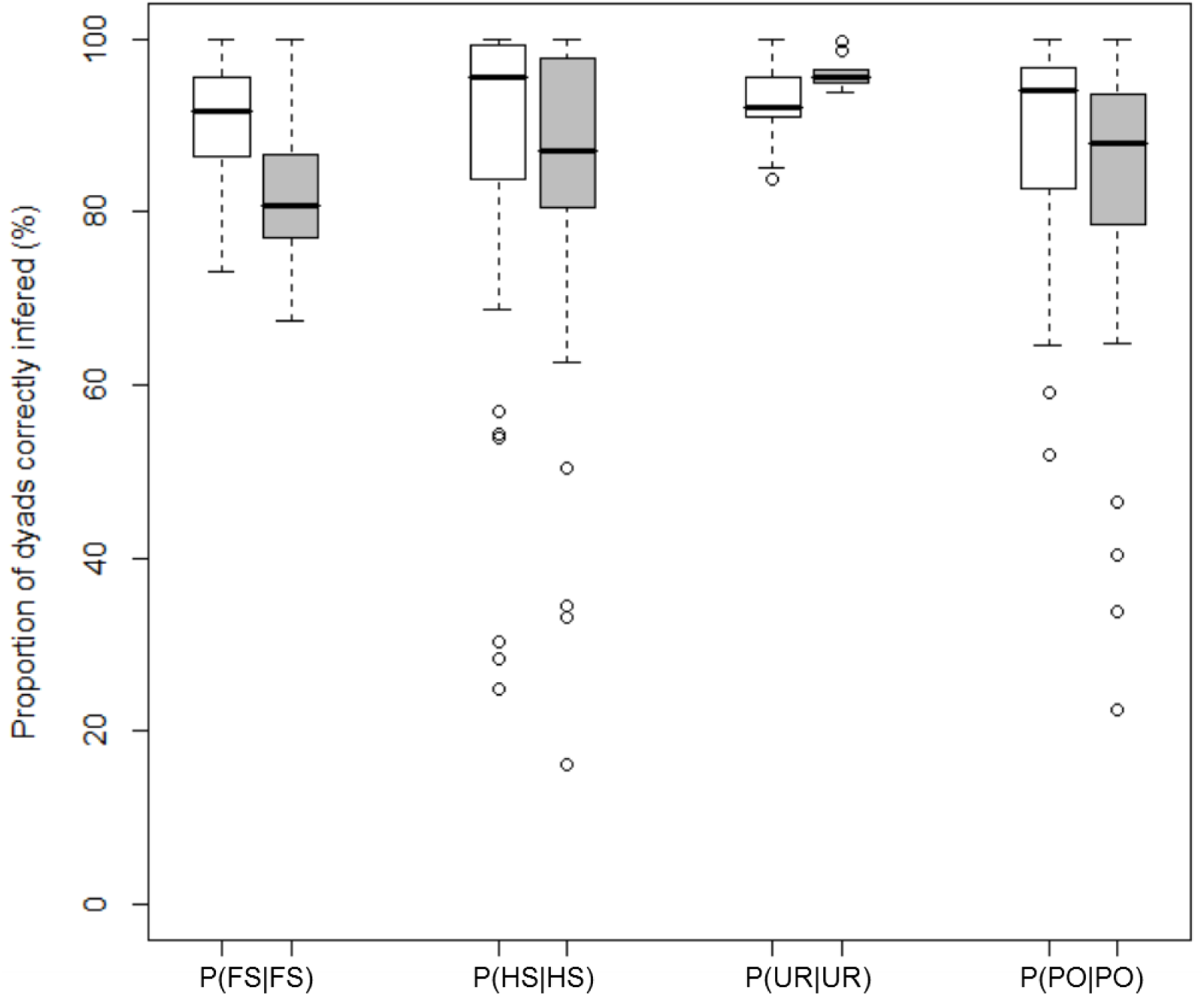
Pedigrees reconstructions from CNV data. The accuracy of pedigrees reconstruction using allele specific copy number (ASCN) genotypes for inherited CNVs only (white boxes) and for inherited and *de novo* CNVs (grey boxes) is estimated with four parameters: the proportion of full-sib [P(FS|FS)], half-sib [P(HS|HS)], unrelated [P(UR|UR)] and parent-offspring [P(PO|PO)] dyads correctly inferred.

### High rates of spontaneous copy number mutations

We analyzed the 3624 trios (corresponding to 7248 meiotic generations) in 54F and 1LF data sets to estimate the rates of spontaneous copy number changes from parent to offspring. The locus specific mutation rate μls at the 113 genes displaying *de novo* CNVs ranged from 2.6 × 10^−4^ to 9.3 × 10^−2^ mutation per generation which is nearly two orders of magnitude broader than the range observed for mammals using the same experimental approach (Table 2). Assuming that *de novo* CNVs have an equal chance of occurring at any location across the gene space, we estimate the mutation rate is 3 × 10^−5^ mutation per gene per generation based on the data obtained for 14,058 genes targeted in our study. This estimate of the cross-genome mutation rate μ_cg_ is one to two orders of magnitude higher than that observed for unicellular and multicellular eukaryotes (Table 2).

Considering the 14058 genes examined here, individuals in 54F and 1LF data sets have inherited non-two-copy number in five genes on average and had 1/6 chance of harboring one to two additional gene(s) with non-two-copy number resulting from spontaneous mutations. Using a poison distribution to predict the number of spontaneous events occurring in the whole gene space (37491 to 56064 genes (De La Torre et al. 2014; Warren et al. 2015)) we estimate a genomic mutation rate u_g_ of 1.4 ± 0.36 per haploid genome per generation which is higher than what was observed in unicellular and multicellular eukaryotes (Table 2) with *P. glauca* u_g_ 3 and 23 times higher than *H. sapiens* and *A. thaliana* u_g_ respectively.

**Table 2:** Copy number mutation rate estimates in *P. glauca* and for other eukaryotes.

| Species | Mutation rate | Reference |
| --- | --- | --- |
| **$\mu_{ls}$ (mutation per generation)** |  |  |
| *P. glauca* | $2.6 \times 10^{-4} - 9.3 \times 10^{-2}$ | This Study |
| *H. sapiens* | $6.5 \times 10^{-3} - 1.2 \times 10^{-2}$ | Itsara et al. 2010 |
| *M. musculus* | $3.6 \times 10^{-3} - 1.1 \times 10^{-2}$ | Egan et al. 2007 |
| **$\mu_{cg}$ (mutation per gene per generation)** |  |  |
| *P. glauca* | $3.0 \times 10^{-5}$ | This Study |
| *S. cerevisiae* | $2.1 \times 10^{-6}$ | Lynch et al. 2008 |
| *M. musculus* | $1.2 \times 10^{-6}$ | Egan et al. 2007 |
| *D. melanogaster* | $9.4 \times 10^{-7}$ | Schrider et al. 2013 |
| *C. elegans* | $2.2 \times 10^{-7}$ | Lipinski et al. 2011 |
| **$u_g$ (mutation per haploid genome per generation)** |  |  |
| *P. glauca* | 1.4 | This Study |
| *H. sapiens* | 0.4 | Sung et al. 2016 |
| *M. musculus* | 0.08 | Sung et al., 2016 |
| *A. thaliana* | 0.06 | Sung et al., 2016 |
| *C. elegans* | 0.04 | Sung et al., 2016 |
| *D. melanogaster* | 0.04 | Sung et al., 2016 |
| *S. cerevisiae* | 0.001 | Sung et al., 2016 |

### Variable CN mutation rates between genes and for different CNV classes

A closer inspection of the mutation rates for different genes revealed a bimodal distribution (mode 1 with μ_ls_ < 10^−2^ and mode 2 with μ_ls_ > 10^−2^ mutation per generation) (Figure 3-A). The μ_ls_ estimates from the large family were more widely spread (Figure 3-B) while estimates from the 54 smaller families covered a narrower range (Figure 3-C). This trend was expected given that the 1LF data set contained around 2000 trees, which should facilitate the detection of rare and recurrent events while the 54F data set may reveal mutation events that are common in the population. Taken together, the estimates from both data sets should provide a more complete picture of the spontaneous mutation dynamics in the *P. glauca* gene space.

**Figure 3:**
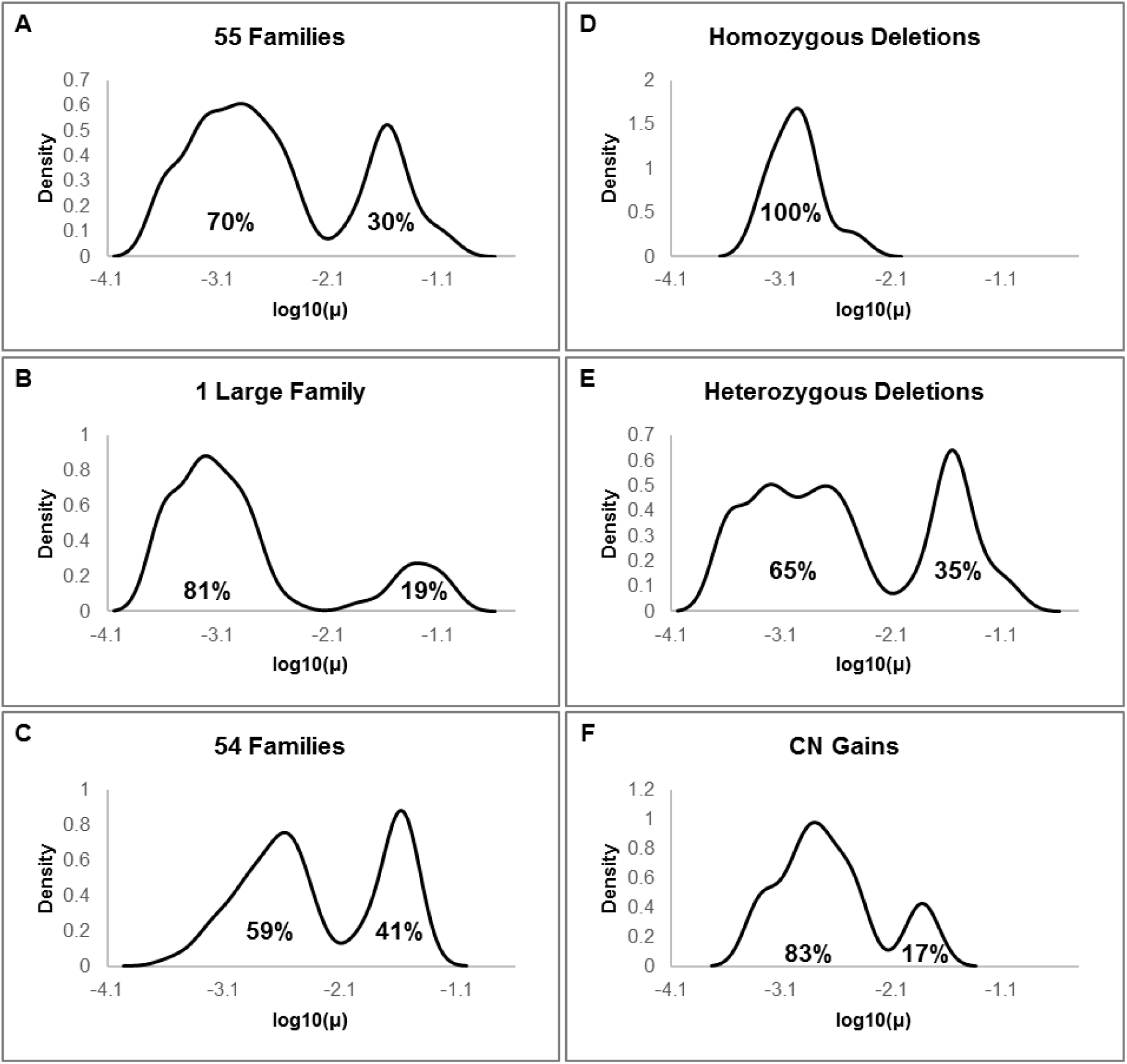
Spontaneous mutation rates distribution. The bimodal distribution of μ_ls_ for all of the families (A), the large family data set (B) and the 54 small families’ data set (C). The distribution of μ_ls_ for different CNV classes: homozygous deletions (D), heterozygous deletions (E) and copy number gains (F). The proportion of CNV genes is shown for each mode (%).

The spontaneous mutation rates varied for different CNV classes. For copy number gains and heterozygous deletions, the mutation rates were mostly in the range of mode 1 with only 17 and 35% of the genes, respectively, in the range of mode 2 (Figure 3-E,F). On the other hand, mutation rates for homozygous deletions were restricted to mode 1 (Figure 3-D).

### Allele specific CN mutation rates

We estimated that allele specific mutation rates μ_AS_ for CNVs were an order of magnitude lower than μ_AS_ for SNPs (Figure 4-A). This observation was based on the analysis of seven genes for which crosses between two homozygote parents allowed us to estimate the μ_AS_ for CNVs and SNPs in the same individuals. Figure 4-B also shows that differences between the mutation rates of two alleles of the same gene Δμ_AS_ can be as high as 10^−2^ mutation per generation for CNVs and SNPs, which has considerable evolutionary consequences.

**Figure 4:**
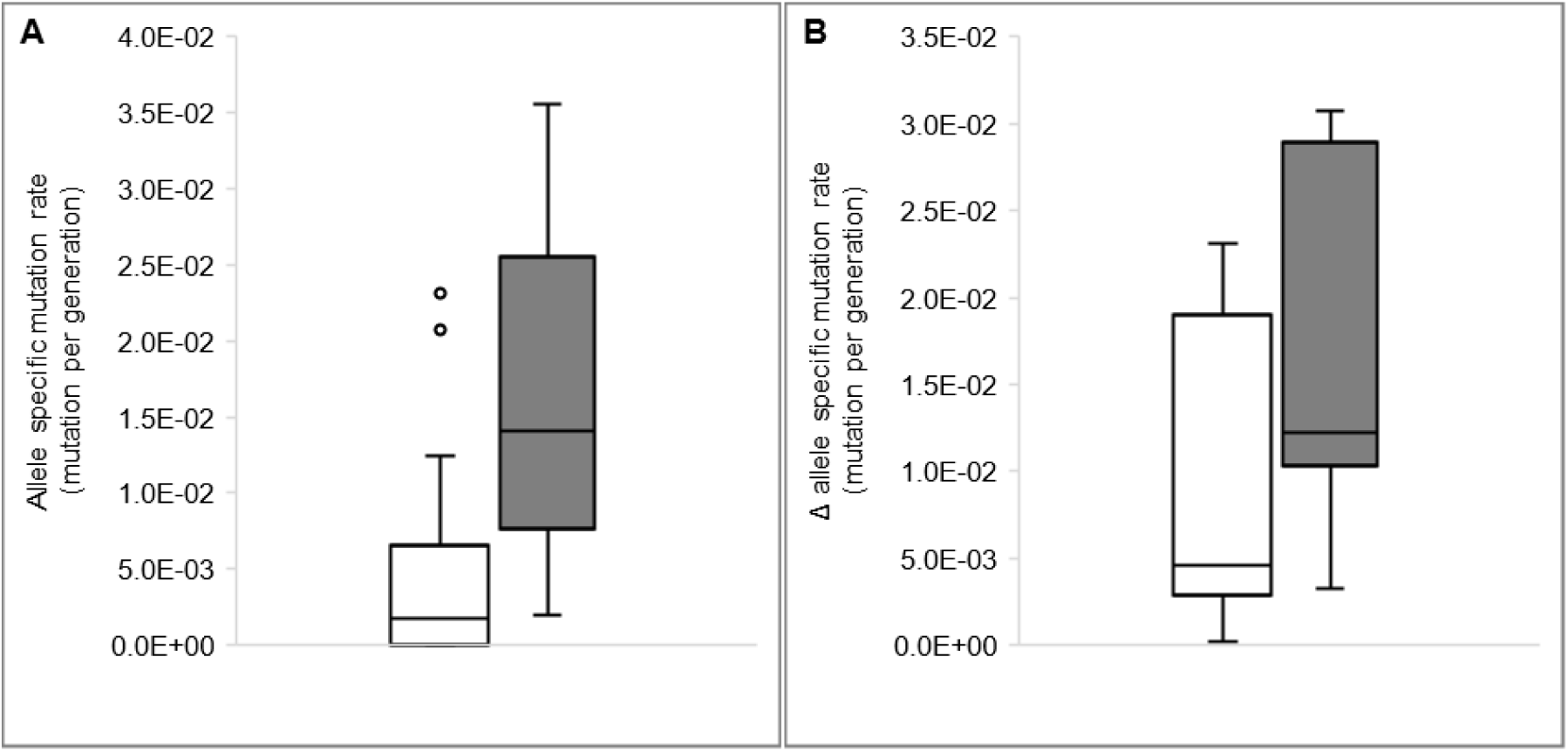
Allele specific mutation rates. Range of the allele specific mutation rates μ_AS_ for the CNVs (white box) and SNPs (grey box) of the 7 loci for which the homozygote individuals were cross-bred (A). Differences between the mutation rates of the two alleles on a locus by locus basis (Δμ_AS_) for CNVs (white box) and SNPs (grey box) variations (B).

### Relationship between CN mutation rates and gene expression

Expression levels for the 113 genes displaying *de novo* CNVs are available for eight *P. glauca* tissues (Raherison et al. 2012) and were analyzed here. We checked if there was an association between the locus specific mutation rate μ_ls_ and transcript accumulation levels. In both data sets (54F and 1LF), we show that the μ_ls_ of mode 2 genes was negatively correlated with gene expression level (Figure 1-A,B) indicating that highly and broadly expressed genes have a lower μ_ls_. This observation does not hold true for mode 1 genes as there was no correlation between μ_ls_ and gene expression levels (Figure 5-C, D).

**Figure 5:**
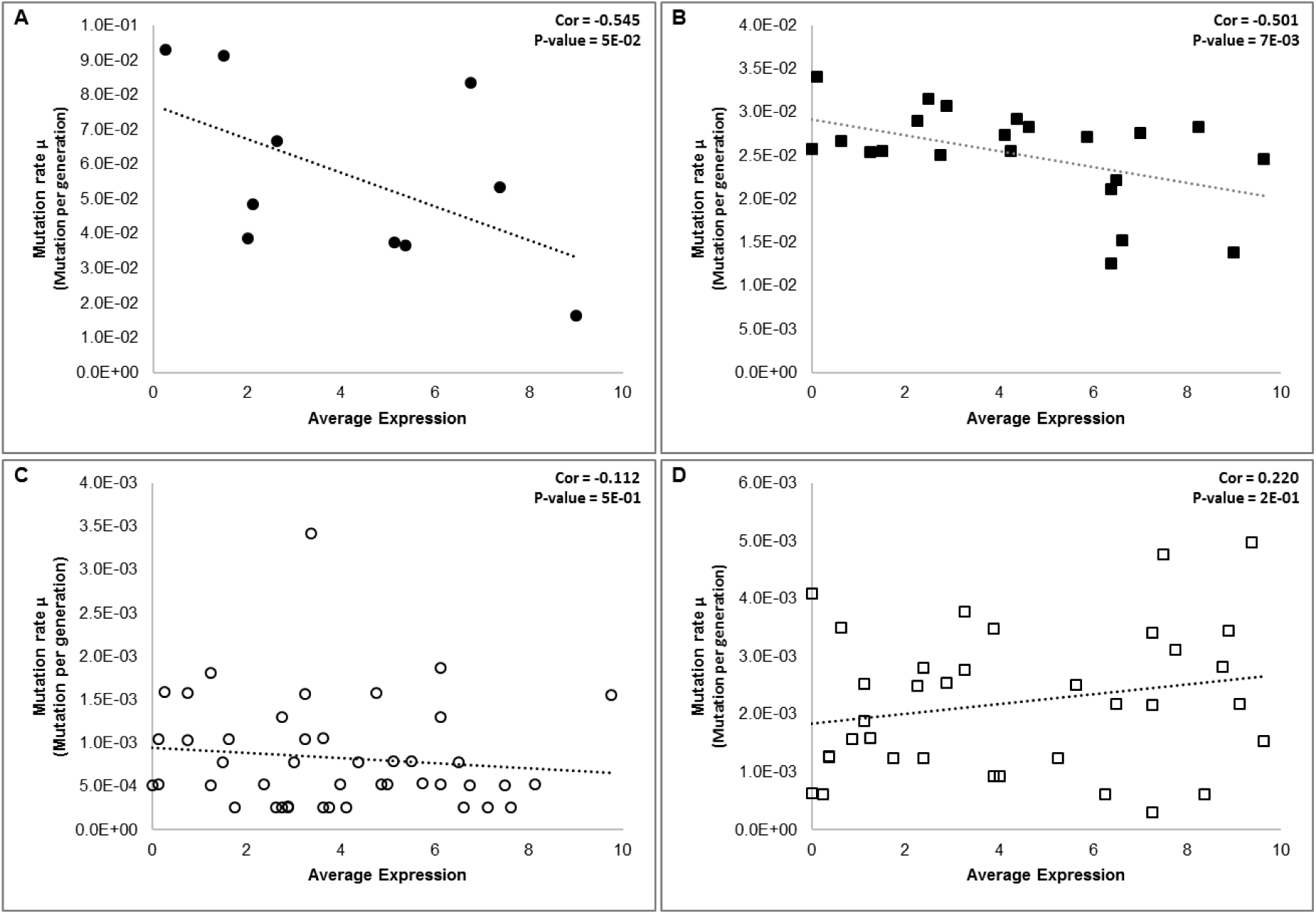
Correlation between mutation rates and gene expression. Negative correlations were identified between the mutation rates and expression levels for mode 2 genes (μ_ls_ > 10^−2^) (A, B) and not for mode 1 genes (μ_ls_ < 10^−2^) (C, D) both for the data sets 1LF (A, C) and 54F (B, D). Average expression was calculated based on the relative transcript accumulation class (1 being lowest and 10 being highest) reported in Raherison et al. (2012).

### CNVs are associated with transmission distortions

We examined the transmission profile of the 23 inherited CNVs detected in the 54F data set for crosses where at least one parent was heterozygote for one copy (A1) and zero copy (A0) alleles. A situation of transmission distortion (TD) is a departure from the expected transmission ratio 0.5 under Mendel’s laws of inheritance. Out of the 23 CNV genes, 16 (70%) displayed transmission ratio distortions (TRDs) while the remaining 7 (30%) were transmitted according to the Mendelian expectations or the number of trios examined was too small to detect TRDs (Table 3). Preferential transmission of one copy (allele A1) was found in 13 (81%) of the TRD genes and preferential transmission of zero copy (allele A0) was found in three (19%) of the TRD genes.

The data revealed that the observed transmission distortions can be dependent on i) whether the heterozygote parent was contributing as male or female (44% of the genes with TRDs show a parental effect: 31% maternal effect and 13% paternal effect) and/or ii) the copy number genotype of the partner in the cross (54% of the genes with TRDs) (Table 3).

**Table 3:** Parental and partner genotype effects on copy number transmission ratio distortion (cnTRD).

| GenBank Acc | ♀He x ♂Ho | ♀Ho x ♂He | ♀He x ♂He | He x N | Favored CN | PE | PGE |
| --- | --- | --- | --- | --- | --- | --- | --- |
| BT103424 | + | - | + | + | 0 | D (ME) | I |
| BT109608 | + | - | - | + | 0 | D (ME) | D |
| BT101196 | - | + | - | - | 0 | I (PME)* | D |
| BT105201 | + | + | NA | + | 1 | I (PME) | ND |
| BT107150 | + | + | + | + | 1 | I (PME) | I |
| CO253730 | + | + | + | + | 1 | I (PME) | I |
| BT110689 | + | + | + | + | 1 | I (PME) | I |
| BT119696 | - | + | NA | + | 1 | D (PE) | ND |
| BT113426 | - | + | - | + | 1 | D (PE) | D |
| BT111788 | + | - | + | + | 1 | D (ME) | I |
| BT106719 | + | - | - | + | 1 | D (ME) | D |
| BT106118 | + | - | - | + | 1 | D (ME) | D |
| DV985927 | - | - | NA | + | 1 | I | ND |
| BT109181 | - | - | - | + | 1 | I | I |
| BT116133 | - | - | + | + | 1 | I | D |
| DR563872 | - | - | + | - | 1 | I | D |
| DR587158 | - | - | NA | - | ND | ND | ND |
| BT103050 | - | - | - | - | ND | ND | ND |
| BT119776 | - | - | NA | - | ND | ND | ND |
| BT101202 | - | - | NA | - | ND | ND | ND |
| BT114401 | - | - | - | - | ND | ND | ND |
| BT110740 | - | - | NA | - | ND | ND | ND |
| BT109138 | - | - | - | - | ND | ND | ND |

\* More details in table 5. He: heterozygote, Ho: homozygote, N: heterozygote or homozygote, CN: copy number allele, PE: parental effect, PGE: partner genotype effect, TRD: transmission ratio distortion, +: significant TRD, -: non-significant TRD, NA: not available, ND: not determined, D: dependent, I: independent, (PE): paternal effect, (ME): maternal effect, (PME): paternal and maternal effects. A more detailed version of this simplified table is presented in Table S1.

We also observed that the level of transmission distortion is dependent on the genetic background of the parents. For a parent participating in different crosses, the level of transmission distortion will vary for different partners (Figure 6-A) as indicated by variable distributions of the transmission ratios TR(A0) for different families (broad, narrow, mono-modal and bi-modal distributions were identified) (Figure S1).

**Figure 6:**
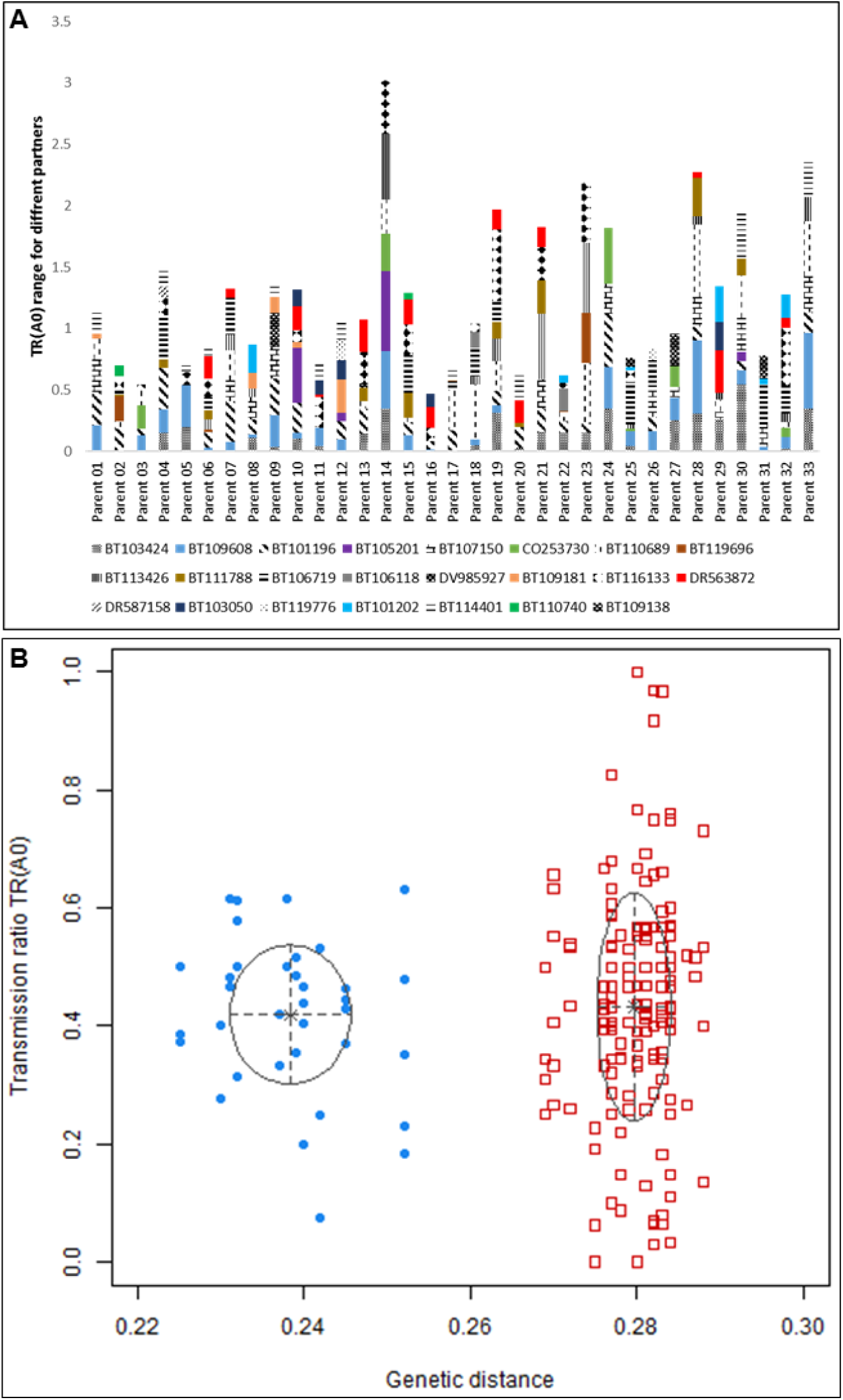
Effects of genetic background and genetic distance between parents on copy number transmission ratio distortion (cnTRD). Transmission ratio range when an individual is crossed with different partners for each of the 23 inherited CNVs (A). Plot of transmission ratios versus the genetic distance between the two parents in the cross (B).

An additional factor that may influence the level of transmission distortion is the genetic distance (gd) between the two individuals in a cross. Our analysis shows that TR(A0) values are distributed into two clusters (Figure 6-B) with average genetic distances of 0.24 and 0.28. The two clusters have the same average distortion level (t-test *p-value* = 6.2E-01) but the TR(A0) variance in cluster 2 (crosses with more distant parents) was twice that of cluster 1, suggesting that crosses involving more distant individuals can give rise to more extreme TRDs. No significant correlations were found between TR(A0) and genetic distance (Cor = -0.20; *p-value* = 2.4E-01 in group 1 and Cor = 0.06; *p-value* = 4.3E-01 in group 2).

### Transmission distortions contribute to CN allele frequency changes between generations

TRDs have the potential to considerably change alleles frequencies on a scale of very short evolutionary periods. Here we show that TRDs can contribute to changes in CN alleles frequencies ranging between 0.001 and 0.08 in a single generation. We also observed that the CN allele frequency change between the parental generation (n) and the next generation (n+1) delta p(A0) was well correlated with transmission ratios TR(A0) (Cor = 0.73; *p-value* = 4.2E-05) (Figure 7-A). That being said, the level of distortion TR(A0) was not the only factor influencing the CN allele frequency changes between generations (Figure 7-B). Chevin and Hospital (2006) proposed a linear model that links the change in allele frequency *Δp* with the transmission ratio *TR* and the product of alleles frequencies in the parental generation *pq*:

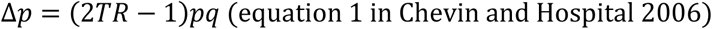

We found that the model fits our data well (adjusted *R^2^* = 0.91; *p-value* = 2.7E-06) for the 13 genes where transmission ratios (TR) are independent of the parental and partner genotype effects. When we considered all of the 23 inherited genic CNVs, the fit (*p-value* = 9.0 E-07) was less strong (adjusted *R^2^* = 0.73) due to the interference of the parental and partner genotype effects on TR(A0) levels for some genes. These results were also confirmed by an ANOVA analysis with *p-value* = 4.0E-06 for the effect of TR(A0) on delta p(A0) and *p-value* = 5.8E-0.4 for the interaction between TR(A0) and pq.

**Figure 7:**
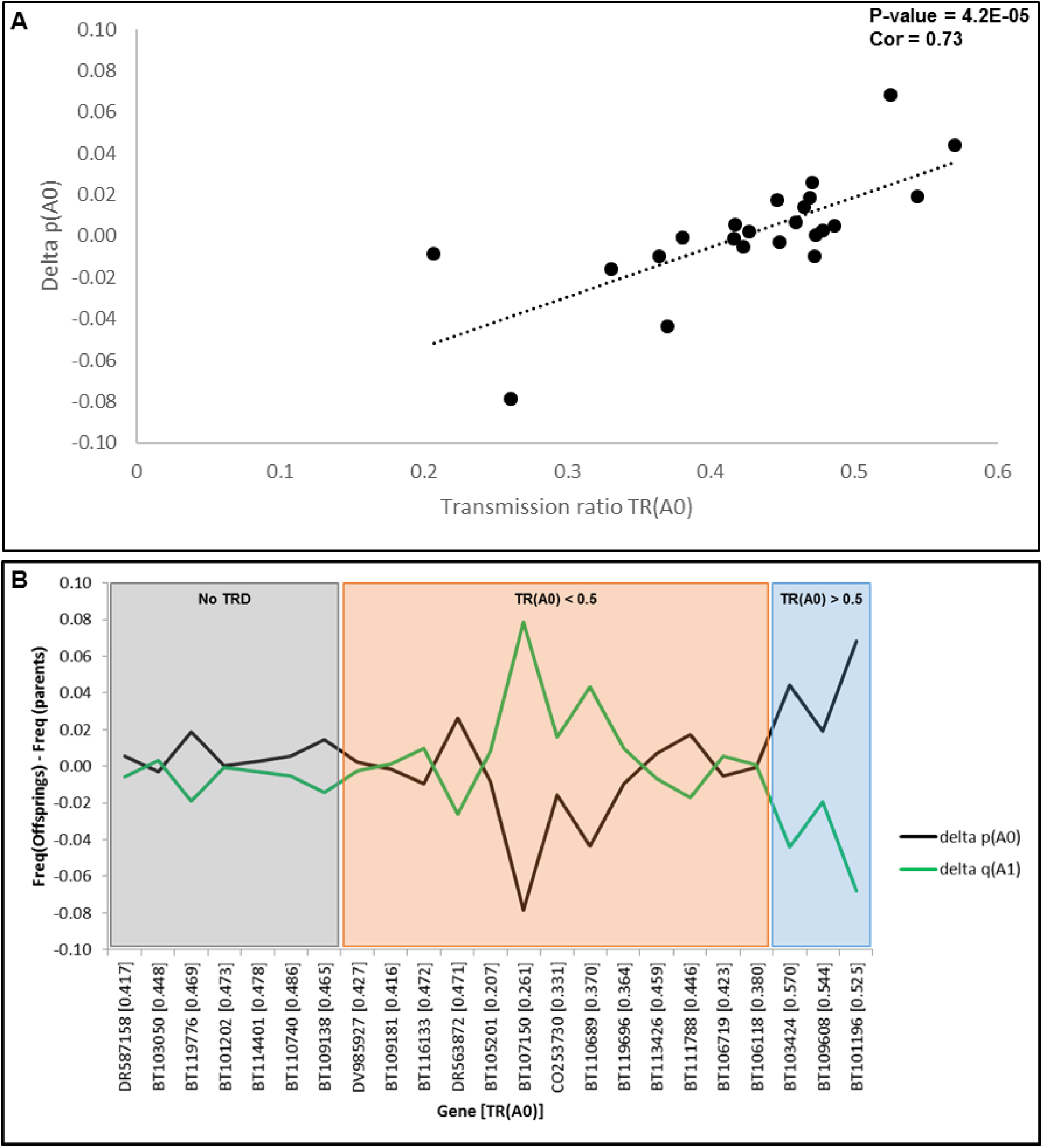
Evolution of copy number alleles frequencies from the parental generation to the offspring generation is function of the transmission ratio TR(A0). Correlation between delta p(A0) (A0 frequency in generation (n+1) – A0 frequency in generation n) and transmission ratio TR(A0) (A). Evolution of A0 et A1 alleles frequencies between generations for the 23 inherited CNVs (B).

### Genes with preferential transmission of zero copy

For the genes (BT101196, BT103424 and BT109608) where TDs favored the allele A0 (zero-copy), we identified three different patterns of selection based on the genotypes frequencies (Table 4). These patterns were inferred from the examination of genotypes and alleles frequencies in the parents and offspring generations in three steps.

First, we considered all the crosses (Table S2) and found that i) for the gene BT101196, there was a significant departure from Mendelian expectations, even if the proportion of crosses with TRD was only (31%), because the level of transmission distortion for this gene was high ii) for the gene BT103424, the level of distortion observed was lower but significant in a pedigree population that included 42% of crosses with TRD and 58% crosses with no TRD (those with two homozygote parents and those where the heterozygote parent was male) and iii) for the gene BT109608, the effect of transmission distortions was diluted in the population because of the interference of both parental and partner genotype effects on TD levels and the presence of crosses between two homozygote parents (crosses with no TRD represent 76% for this gene).

In the second step, we considered only the crosses with at least one heterozygote parent (Table S3) and observed that for the three genes analyzed, there was a significant departure from the genotypes frequencies expected in the case of a Mendelian inheritance, although the detected level of distortion was moderate due to the interference of the parental and partner genotype effects that still remained in the pedigree population.

Finally, we examined only the crosses with significant TDs (Table S4) and were able to quantify the effect of transmission distortions on changes in genotypes frequencies between generations without the interference of double homozygote crosses, parental or partner genotype effects. For the three genes, there were significant deviations from the expected genotypes frequencies and the levels of distortion were higher than those observed in the second step of the analysis.

The findings of this analysis are summarized in Table 4 and show that for the gene BT101196 (highly similar to F-box proteins), the frequency of the allele zero copy (A0) increased in the offspring resulting from more of the one copy (A0/A1) genotype and a lower frequency of the two copy genotype (A1/A1), which suggests that it is under balancing selection with a heterozygote advantage. On the other hand, for the two other genes BT103424 (unknown function) and BT109608 (Chaperone DnaJ-domain superfamily protein), the increased frequency of the allele A0 was due to more of the zero copy (A0/A0) genotype, which suggests that they are under directional selection favoring the allele (A0) and the homozygote genotype (A0/A0) at the expense of the genotype (A1/A1) or both the genotypes (A0/A1) and (A1/A1) for the genes BT103424 and BT109608, respectively.

**Table 4:** Three patterns of selection on the CN genotypes of genes favoring the transmission of zero copy.

| Gene | BT101196 | BT103424 | BT109608 |
| --- | --- | --- | --- |
| **Predicted function** | F-box protein | Unknown | Chaperone DnaJ-domain protein |
| **Selection pressure** | Balancing selection | Directional selection | Directional selection |
| **Selection pattern** | 1 | 2 | 3 |
| **A0/A0 genotype (Zero copy)** | Neutral | Advantageous | Advantageous |
| **A0/A1 genotype (One copy)** | Advantageous | Neutral | Deleterious |
| **A1/A1 genotype (Two copies)** | Deleterious | Deleterious | Deleterious |

### The particular case of an F-box gene

The gene BT101196 attracted our attention because it is homologous to an *A. thaliana* gene involved in embryo development arrest (Pagnussat et al. 2005) and both the *P. glauca* and *A. thaliana* genes display interesting but distinct transmission distortions. In *P. glauca*, the transmission of the zero copy allele is favored (TR(A0) > 0.5) and is influenced by the genotype of the partner. We observed that transmission distortions occur only when the heterozygote parent is crossed with an individual harboring two copies of the gene, i.e. no distortion is observed when the partner has a zero or one copy genotype (Table 5). The data also show simultaneous paternal and maternal effects on TR(A0) with transmission distortions being observed whether the heterozygote parent contribute as male or female (Table 5). Like the other genes displaying TRDs, the gene BT101196 transmission was dependent on the parents’ genetic background but not the genetic distance between them (Figures S2 and S3).

**Table 5:** Parental and partner genotype effects on copy number transmission ratio distortion (cnTRD) for the F-box gene BT101196.

| **Partner genotype effect** |  |  |  |  |
| --- | --- | --- | --- | --- |
| **Crosses** | **N** | **TR(A0)** | **CI (95%)** | **P-value** |
| (A0/A1) x (A1/A1) | 458 | 0.598 | 0.552 – 0.643 | 3.0E-05 |
| (A0/A1) x (A0/A1) | 244 | 0.488 | 0.423 – 0.552 | 7.5E-01 |
| (A0/A1) x (A0/A0) | 402 | 0.494 | 0.459 – 0.529 | 7.5E-01 |
| **Maternal effect** |  |  |  |  |
| **Crosses** | **N** | **TR(A0)** | **CI (95%)** | **P-value** |
| ♀ (A0/A1) x ♂ (A1/A1) | 239 | 0.577 | 0.512 – 0.641 | 2.0E-02 |
| ♀ (A0/A1) x ♂ (A0/A0) | 168 | 0.476 | 0.399 – 0.555 | 5.9E-01 |
| **Paternal effect** |  |  |  |  |
| **Crosses** | **N** | **TR(A0)** | **CI (95%)** | **P-value** |
| ♀ (A1/A1) x ♂ (A0/A1) | 219 | 0.621 | 0.553 – 0.686 | 4.2E-04 |
| ♀ (A0/A0) x ♂ (A0/A1) | 76 | 0.513 | 0.396 – 0.630 | 9.1E-01 |

A main difference in the transmission distortion of the *P. glauca* gene BT101196 and its *A. thaliana* homolog (MEE66, AT2G02240) is that A0 was favored in *P. glauca* and A1 was favored in *A. thaliana* (Pagnussat et al. 2005). This and other TRD features, along with gene expression and embryo viability phenotypes, are summarized in Table 6.

**Table 6:**
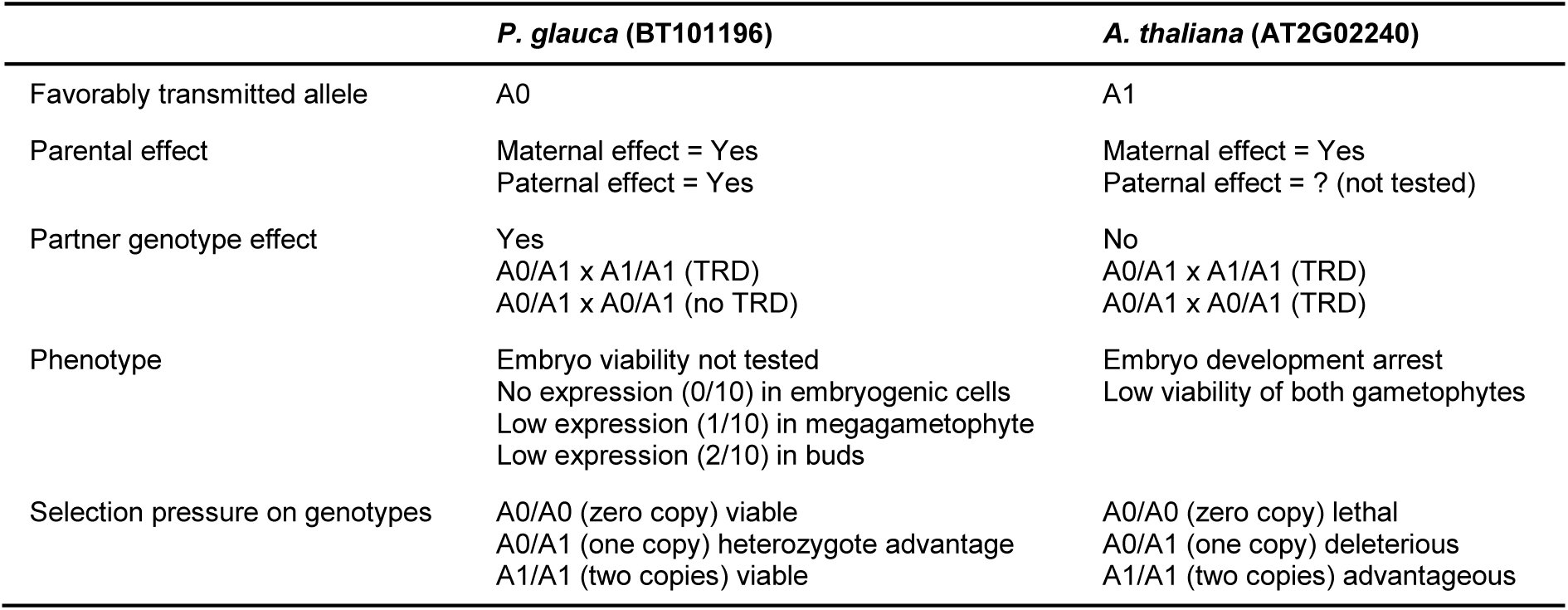
F-box gene copy number transmission in *P. glauca* and *A. thaliana*.

|  | *P. glauca* (BT101196) | *A. thaliana* (AT2G02240) |
| --- | --- | --- |
| Favorably transmitted allele | A0 | A1 |
| Parental effect | Maternal effect = Yes<br>Paternal effect = Yes | Maternal effect = Yes<br>Paternal effect = ? (not tested) |
| Partner genotype effect | Yes<br>A0/A1 x A1/A1 (TRD)<br>A0/A1 x A0/A1 (no TRD) | No<br>A0/A1 x A1/A1 (TRD)<br>A0/A1 x A0/A1 (TRD) |
| Phenotype | Embryo viability not tested<br>No expression (0/10) in embryogenic cells<br>Low expression (1/10) in megagametophyte<br>Low expression (2/10) in buds | Embryo development arrest<br>Low viability of both gametophytes |
| Selection pressure on genotypes | A0/A0 (zero copy) viable<br>A0/A1 (one copy) heterozygote advantage<br>A1/A1 (two copies) viable | A0/A0 (zero copy) lethal<br>A0/A1 (one copy) deleterious<br>A1/A1 (two copies) advantageous |

## Discussion

In this work, we took advantage of the availability of raw intensity data from a SNP-array for several thousand genes obtained from thousands of individuals to identify and characterize genic CNVs. We applied a family-based approach coupled with a cross-samples strategy (for CNV calls) that allowed us to detect high-quality copy number genotypes. We obtained new insights into the contribution of different evolutionary forces (spontaneous mutations, transmission distortions, selection and random genetic drift) to the generation, transmission and maintenance of a class of genetic variations, which have been poorly studied in the genome of perennial plants. The approach is transferable to other species, including non-model organisms and species with large and complex genomes or long generation times.

### CNVs in the *P. glauca* gene space

CNVs can affect gene structure and expression, and impact downstream phenotypes, fitness and reproductive success (Tang and Amon 2013). Consequently, many CNVs are expected to be deleterious and under strong purifying selection. Our data showed that only 0.5 to 1% of the 14058 targeted genes displayed CNVs even though thousands of individuals were examined. Based on our data, we predict that each individual will have on average 17 to 20 genes (0.04% of the gene space) with non-two-copy genotypes. The majority of CNVs identified in this study were copy number losses (90%), and bi-allelic variations were more abundant by far than multi-allelic variations, as previously observed in other species including human (Mills et al. 2011; Kato et al. 2010), stickleback fish (Chain et al. 2014), bovine (Cicconardi et al. 2013) and plants (Swanson-wagner et al. 2010; Yu et al. 2011; McHale et al. 2012), to cite few. These observations agree with the afore mentioned hypothesis of purifying selection. In our CNV set, copy number losses were nine times more abundant than copy number gains. This estimate is similar to ratios of seven and height reported for maize by Swanson-wagner et al. (2010) and Liu et al. (2012), respectively. Rice (Yu et al. 2011) and soybean (McHale et al. 2012) also showed a bias toward copy number losses. In human however, the ratio of deletions to duplications is considerably lower (two to three times) (Chen et al. 2010) than in plants. This detection of more abundant copy number losses can be explained from technical and/or biological perspectives. For large copy numbers, signal intensities become noisy and the relationship between copy number and signal intensity is not linear (Cantsilieris, Baird, and White 2013) which makes the detection of copy number gains less likely. Also, multiple copies of a gene can harbor sequence polymorphisms that may cause mismatches during probe and primer hybridizations. From a biological perspective, several molecular mechanisms involved in CNV formation favor sequence losses over duplications (Chen et al. 2010), particularly the non-allelic homologous recombination (NAHR) mechanism shown to be the dominant mechanism for CNV formation in *A. thaliana* (Lu et al. 2012). Another, non-exclusive, hypothesis is that losses are the result of the segregation of non-allelic homologs. In maize, single-copy homologous sequences located in non-allelic positions in the genomes of two crossed hemizygous parents, segregate in the offspring as zero, one or two copies genotypes (Liu et al. 2012), which is consistent with our observations for most of the genes in the present set of *P. glauca* families.

### Copy number spontaneous mutations represent a non-negligible evolutionary force

To date, estimates of the mutation rate for CNVs have been limited to a few model organisms and suffer from biases related to the targeted region in the genome, the individuals sampled (families or populations), and the estimation approach (Katju and Bergthorsson 2013; Itsara et al. 2010). Therefore, a better characterization of the CN mutation rates spectrum in different species is expected to provide new insights into the evolution process.

Here we show that CN mutation rates in *P. glauca* cover a wide range (three orders of magnitude) and can reach as high as 10^−2^ mutation per generation for some genes. High spontaneous mutation rates (locus specific and genomic estimates) are expected in plants and particularly trees. In plants, many cell divisions occur during a single generation, which increase the probability of mutation events during DNA replication and repair (Petit and Hampe 2006; Scofield and Schultz 2006). More importantly, there is no clear separation between germline and soma which both contribute to the estimated mutation rates in plants. Somatic mutations are particularly high in plants and are frequently generated under stress; for example, a two-fold increase in μ was found for stress induced mutations (Jiang et al. 2014; Debolt 2010). Mutations generated during plant growth, accumulate in the meristem and are transmitted to the gametes. This phenomenon may be more pronounced in perennials (like *P. glauca*) compared to annuals due to their longevity. In conifers, the egg cell for fertilization differentiates from the megagametophyte through about 11 rounds of mitotic division from the megaspore and the sperm nucleus is formed via five mitotic divisions from the microspore (Williams 2009). These rounds of division between meiosis generating the megaspore or the microspore, and the fecundation increase the chances of spontaneous mutations. The high mutation rates reported here can also be explained by two particular features of the *P. glauca* genome: a high A-T content (62%) and an abundance of repeated sequences (70%), particularly transposons (Birol et al. 2013; Nystedt et al. 2013). In other species, transposons were found to actively promote high mutation rates (Woodruff et al. 1984; Bégin and Schoen 2006; Lu et al. 2012; Pinosio et al. 2016), and the examination of deletions breakpoints identified the presence of A-T rich sequences in the vicinity of these variations (Chen et al. 2010).

Copy number mutation rates in the *P. glauca* gene space followed a bimodal distribution with the majority of genes (70%) subject to low mutation rates and the rest (30%) associated with high mutation rates (above 10^−2^ mutation per generation). This spectrum of mutation rates could reflect local differences in the genome or differences in the selection pressure. Local features of genome architecture such as base composition, short repeats density, mobile elements, recombination rates and methylation can influence the frequency at witch mutations are generated (reviewed in Baer et al. 2007). Also genes involved in basic metabolic functions are expected to be under strong selection pressure. On the other hand, redundant genes, genes associated with compensation mechanisms and genes involved in adaptation are likely to be under relaxed selection and tolerate more frequent mutations (Tang and Amon 2013). In this work, we have shown that homozygous deletions (complete gene losses) are rare (confined in mode 1 only). Since the complete loss of a gene is expected to be more deleterious than a partial loss (heterozygous deletion) or a duplication, we can presume that these mutations are under strong purifying selection.

We found a negative correlation between mutation rate and average gene expression level only for the genes at the high end of the mutation rate spectrum (mode 2). This was taken as an indication that selection pressure maintains the mutation rate at lower level for highly and broadly expressed genes which are presumed to be more essential for cellular function. In eukaryotes, the relationship between genes transcription levels and the mutation rates is still not clear. The transcription-coupled repair hypothesis TCRH (Fidantsef and Britt 2011; Hendriks et al. 2010) suggests that highly expressed genes should be associated with lower mutation rates based on the observation that DNA repair is more efficient for actively expressed genes and for the transcribed strand rather than the non-transcribed strand. On the other hand, the transcription-associated mutagenesis hypothesis TAMH (Park, Qian, and Zhang 2012; Sollier et al. 2014; Heinäniemi et al. 2016) proposes that transcription promotes spontaneous mutations based on the observation that highly expressed genes are more frequently associated with mutations (SNPs, intra-genic deletions or double strand breakages). Since we observed a negative correlation only in mode 2, neither the TCRH nor the TAMH hypotheses explain our data entirely. Alternatively, Lynch (2011) proposed the drift-barrier hypothesis DBH, which stipulates that selection will drive down μ from high mutation rates to lower levels while at the lower bound of observed mutation rates, genetic drift will circumvent selection and the mutation rate will evolve randomly toward higher or lower values. This DBH fits our data and explains the observed relationship between CN mutation rate and gene expression in both mode 1 and mode 2.

We found that CNVs have lower mutation rates (an order of magnitude on average) than SNPs for the same genes. In *A. thaliana* and human, the mutation rate for SNPs is one and three order(s) of magnitude higher than for CNVs respectively (Ossowski et al. 2010; Itsara et al. 2010). The effects of CNVs on gene structure and/or expression are likely to be more detrimental on average than those of single base mutations (for example synonymous SNPs are mostly slightly deleterious or neutral). Hence, a strong selection pressure is expected to drive CNV mutation rates to lower levels.

Mutations are the source of variations that fuel the evolutionary process but because mutation events are rare, their contribution to the changes of alleles frequencies and to the determination of evolutionary outcomes has been underestimated. Yampolsky and Stoltzfus (2001) proposed an origin-fixation model in which mutation rates can be an orienting factor in evolution. In this model the fate of two alleles is not determined by their relative effects on fitness alone but also depends on the order of the appearance of alleles, their respective mutation rates and the population effective size (Ne). Here we report empirical estimates of the differences between the mutation rates associated with two alleles of the same locus. A pair of alleles may have mutation rates that differ by as much as an order of magnitude in favor of one allele relatively to the other, which will have a large impact on the chances of fixation of the two alleles. The mutation rate also determines how long an allele will remain in the genome. Alleles with lower deletion rates will be retained longer increasing their chance of accumulating more mutations or being converted to alternative allelic forms.

Copy number mutation rates in *P. glauca* can reach high values, are variable for different genes, alleles and CNV classes and are subject to a variety of selection pressures. These features undoubtedly, make spontaneous copy number mutations a significant orienting factor in evolution and in shaping genome architecture, which in turn determine the fate of genetic variants and of the individuals harboring them.

### Copy number transmission distortions contribute to considerable changes in alleles frequencies between generations

Alleles are maintained in a population according to various factors including their effect on the fitness and reproductive success of the individuals harboring them. Transmission distortions (TDs) may circumvent selection pressure and promote the transmission of an allele, even if it is deleterious to the organism. TD is supposed to be a transient state that will lead to a rapid fixation of the favored allele unless antagonistic forces intervene to maintain allelic polymorphism (Taylor and Ingvarsson 2003). TDs are very common in plants. In *A. lyrata* 50% of the inspected loci displayed transmission ratio distortions (TRDs) (Kuittinen et al. 2004). In conifers, depending on the species, 2 to 79% (with an average of 20%) of the loci examined by Krutovskii et al. (1998) were associated with transmission distortions. Our analysis identified transmission distortions for 70% of inherited CNVs in *P. glauca*, which is at the upper end of the range reported by Krutovskii et al. (1998) for conifers. Out of the 16 CNV genes with TRDs, the majority (81%) favor the transmission of one-copy allele (A1) instead of zero-copy allele (A0) which helps to maintain two-copy genotypes and counteracts the accumulation of copy number losses by drift and mutations.

TRD levels reported in other species are between 0.3 and 0.6 except in a few cases (reviewed in Huang et al. 2013). In our study, TRD levels for preferential transmission of one-copy (A1) and zero-copy (A0) were 0.06 – 1.00 and 0.34 – 1.00, respectively, for different families. For the aggregated data, these ranges are 0.53 – 0.79 and 0.53 – 0.57, respectively. These observations show that even within one species, TRD levels vary widely and can be as extreme as to systematically transmit an allele at the expense of the other (for TR approaching 0 or 1). Allele frequency changes contributed by TRDs for some *P. glauca* CNV genes can reach 0.08 within one generation (present study), which predicts that the favored allele could reach fixation in less than 10 generations. This indicates that transmission distortion may be a strong evolutionary force capable of shaping the genetic diversity in a short evolutionary period.

TRDs can result from mechanisms operating during the gametic (pre- or post- meiosis) or the zygotic stage (pre- or post- fecundation). TRDs can also be parent-sex dependent sdTRDs (paternal effect only or maternal effect only) or independent siTRDs (both, paternal and maternal effects). sdTRDs have a gametic origin and are frequent at the intra-population level. On the other hand, siTRDs have a zygotic origin, play a large role at the inter-population level, are rare within a species and display higher distortion levels than sdTRDs (Huang, Labbe, and Infante-Rivard 2013; J Leppälä et al. 2008; Yohei Koide et al. 2008; Y Koide et al. 2012). The majority of TRDs reported in *A. lyrata* (Leppälä et al. 2013) are sdTRDs and in *A. thaliana* 46% of the identified TRDs are sex-independent (Pagnussat et al. 2005). In *P. glauca*, we found 42% siTRDs and these siTRDs are associated with higher levels of distortion compared to sdTRDs for TRDs favoring the transmission of one copy. For TRDs favoring the transmission of zero-copy, sdTRDs levels were slightly higher than siTRDs, although the number of observations was too small to draw strong conclusions.

Transmission distortion is a genetically controlled mechanism that can be influenced by the partner genotype at the TRD locus, as well as by the genetic background and the genetic distance between the partners. Incompatibilities between alleles of the TRD locus (locus S in *A. lyrata* (J Leppälä et al. 2008) and locus D in monkeyflower (Fishman and Willis 2005)) or of linked and/or unlinked loci located elsewhere in the genome (e.g. in wheat, Friebe et al. (2003); and in rice, Koide et al. (2012)) can influence TRD levels. In this study, we show that a considerable proportion of TRDs (54%) was influenced by the genotype of the partner at the TRD locus. Also, consistent with observations in other plants (Koide et al. 2012; Johanna Leppälä, Bokma, and Savolainen 2013; Buckler et al. 1999), we found that TRDs levels vary considerably for different genetic backgrounds in the parents. This dependence of TRD levels on the genetic background can be the result of different non-exclusive mechanisms: i) the action of a linked or unlinked driver (distorter or suppressor), ii) the epistatic interactions between different loci in the genome or iii) the effects of cytoplasmic factors. We were unable to identify which mechanism is implicated in the genetic control of TRDs in *P. glauca* because we lacked information on the co-segregation of TRD loci with other genomic markers in reciprocal crosses. It was suggested that transmission distortions contribute to the establishment of reproductive barriers and TRD levels should increase linearly (Johanna Leppälä, Bokma, and Savolainen 2013) or exponentially (snow-ball effect) (Moyle and Nakazato 2010) with the genetic distance between parents and the divergence between populations. So far though, the existence of this relationship is controversial because it was detected in some species (Matsubara et al. 2011; Koide et al. 2008; Johanna Leppälä, Bokma, and Savolainen 2013) but not in others (Johanna Leppälä, Bokma, and Savolainen 2013). Our analysis shows no significant association between TRD levels and genetic distance between parents in *P. glauca*. However, TRD levels were more variable with higher genetic distances; therefore, the influence of genetic distance on TRD level could manifest itself in more diverged populations.

Three cases where heterozygote parents preferentially transmitted a zero-copy allele were identified in *P. glauca*. The genes involved likely responded to different selection pressures that promoted their partial or complete suppression from the population. The transmission of the BT103424 (Unknown protein) and BT109608 (Chaperone DnaJ-domain protein) genes was subject to a maternal effect (and most likely have a gametic origin). The one-copy allele (A1) of both genes was under negative selection while the zero-copy allele (A0) was under positive selection (accumulating in the form of the homozygote genotype A0/A0 at the expense of the heterozygote genotype A0/A1 and/or homozygote genotype A1/A1). On the other hand, the TRD for the gene BT101196 (F-box protein) was sex-independent (with both paternal and maternal effects), dependent on the partner genotype (TRD observable only when a heterozygote is crossed with an individual with two copies of the gene) and most likely had a zygotic origin. Its zero-copy (A0) and one-copy (A1) alleles appear to be under balancing selection with a heterozygote advantage. The BT101196 transcripts were not detected in the embryo but were expressed at low levels in the megagametophyte and vegetative buds (Raherison et al. 2012). This pattern suggests a case of sheltered load were the alleles A0 et A1, which have opposing effects on the fitness at different life stages, are both maintained in the population in the form of heterozygote genotypes. The genotype frequencies observed for the gene BT101196 suggest that the allele A1 is likely deleterious (but not lethal) for the embryo and its expression indicates it is necessary in vegetative buds later in development. In *A. thaliana*, transmission of its homolog AT2G02240 (designated by MEE66) is also distorted under a maternal effect, but in contrast to *P. glauca*, the allele A1 is preferentially transmitted to the next generation (Pagnussat et al. 2005) and is under positive selection (Table 6). The different transmission behaviors of these F-box genes, in the two species, likely reflects the different evolutionary processes operating in annual angiosperms and perennial gymnosperms.

### Evolutionary consequences of copy number spontaneous mutations and transmission distortions

In the present study, we primarily detected copy number losses and estimated that CNVs affect a small proportion of the gene space, which supports the hypothesis that CNVs are mainly deleterious and should be under strong purifying selection. We also show that CNVs can be either inherited from the parents or generated via spontaneous mutations. *De novo* CNV formation occurs at a lower rate than SNP formation (expected to be less deleterious than structural variations on average), particularly for complete gene losses (homozygous deletions). Still, copy number mutation rates can reach high levels for *P. glauca* and the genomic mutation rate for this species is higher than in other organisms examined to date. This high mutation rate imposes a mutational load that seems to be tolerated because frequent mutations would fuel the standing genetic variation of the population, contributing to adaptation to environmental changes, which is a well-known feature for perennial trees. Our data further show that alleles of the same gene can have considerably different mutation rates and consequently the fate of alleles can be determined based on the order and rate of their generation and maintenance in the genome in addition to their respective effects on fitness. From this perspective, spontaneous mutations are not only a source of new variants but play also a role as an orienting factor that can determine the fate of alleles.

Our results show that an individual may harbor non-two-copy genotypes in different loci due to inherited copy number variants at least five times more than variants generated *de novo*. However, the transmission of CNVs from a generation to the next is distorted most of the time. The majority of the observed transmission distortions favor the transmission of the one-copy allele and the restoration of the two-copy genotype in the next generation. This pattern is expected in order to maintain the integrity and stability of the diploid genome. On rare occasions, transmission distortions would promote the inheritance of the zero-copy allele either because this variant would behave selfishly or the copy number loss would be advantageous (variant under balancing or positive selection). The present study shows that transmission distortions can cause large allele frequency changes on short evolutionary periods, and that the level of distortion is genetically controlled, which contributes to the maintenance of a substantial standing genetic variation in the population.

## Acknowledgments

We thank Sylvie Blais from the Canada Research Chair in Forest and Environmental Genomics at Université Laval (Quebec, Canada) for help with data manipulation. We also thank Julien Prunier and Mebarek Lamara from Université Laval for help with the collection of samples. We thank the Ministry of Forests, Wildlife and Parcs (Quebec, Canada) for the maintenance of the experimental plantations where the samples were collected. We are thankful to Cyril Van Ghelder from the University of Oxford (Oxford, United Kingdom) for his help with the *in silico* prediction of the domains of the gene BT102213.

Funding was received from Genome Canada and Genome Quebec for the SMarTForests project. AS received financial support from the SMarTforests project and from Quebec Ministry of Economic Development, Innovation and Exports through the GenAC project.

### Competing interests

The authors declare that no competing interests exists.

## Supplemental material

**Figure S1:**
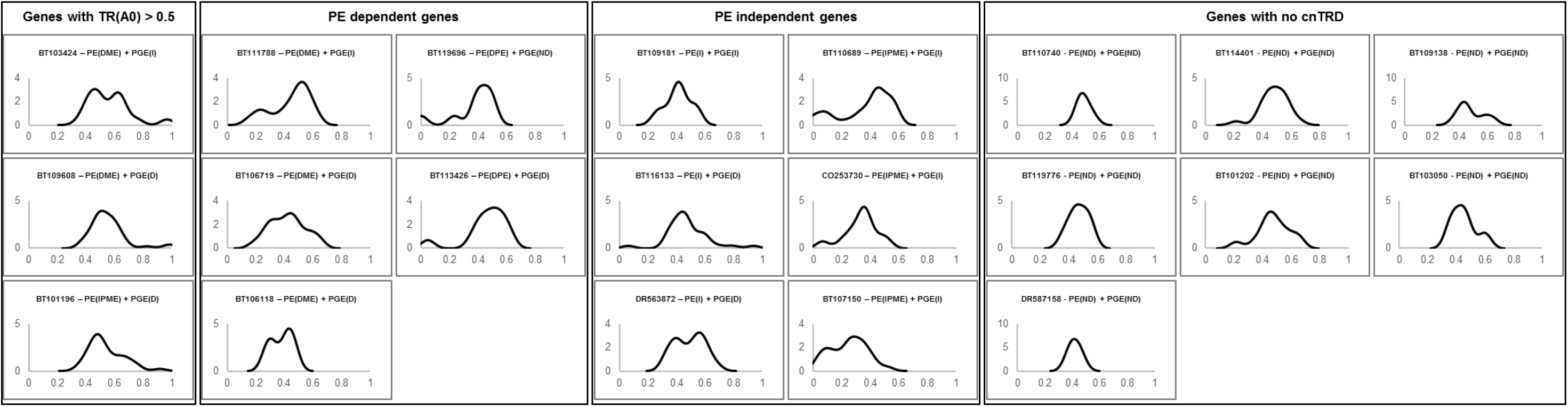
Genetic background effect on copy number transmission ratio distortion (cnTRD). The distribution of transmission ratio TR(A0) for different families. x-axis: TR(A0) values, y-axis: density, PE: parental effect, PGE: partner genotype effect, I: independent, D: dependent, IPME: Independent with both paternal and maternal effects, DPE: dependent with paternal effect only, DME: dependent with maternal effect only, ND: not determined.

**Figure S2:**
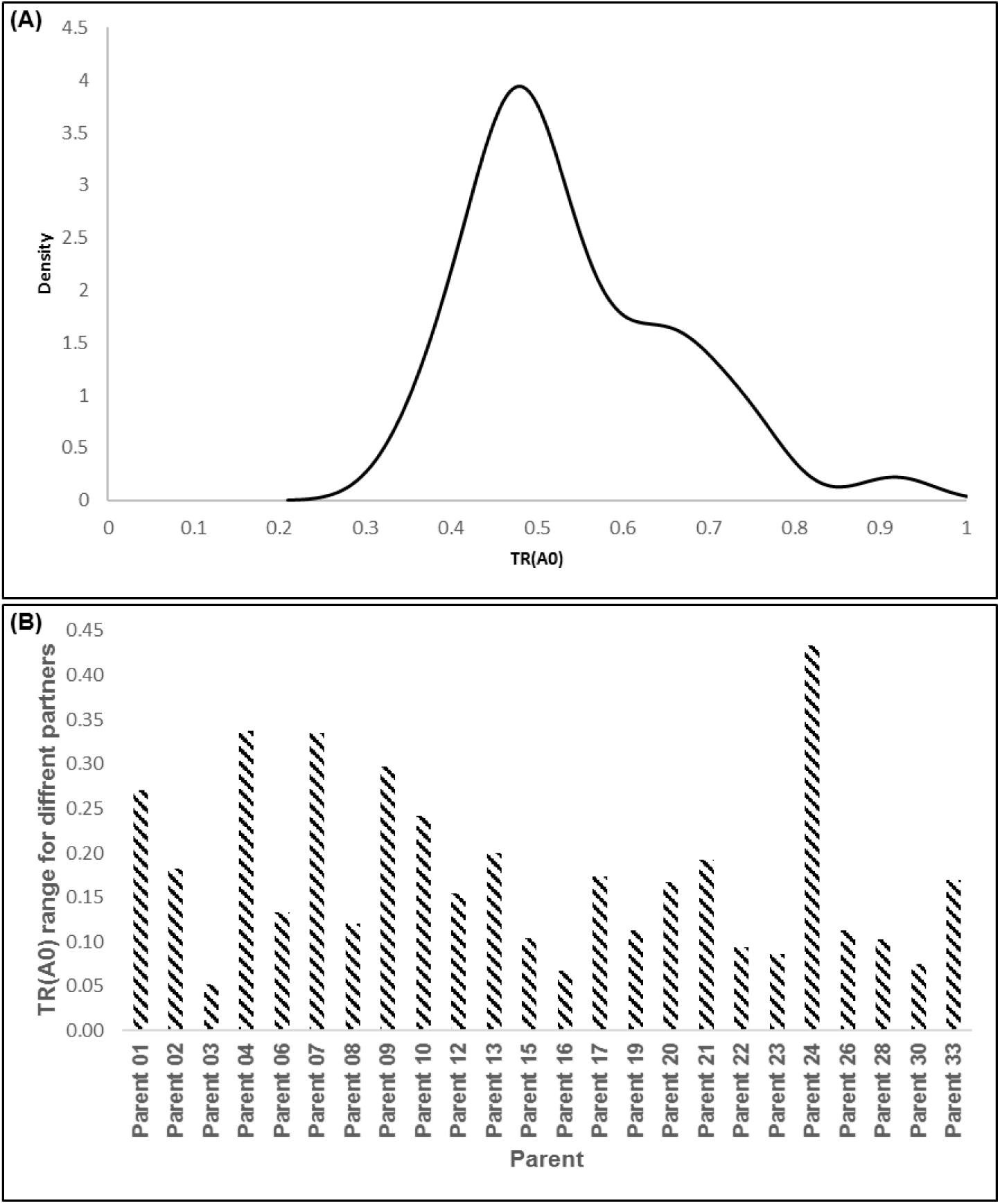
Genetic background effect on copy number transmission ratio distortion (cnTRD) for the F-box gene BT101196. Distribution of transmission ratio TR(A0) for different families (A). Transmission ratio range when a parent is crossed with different partners (B).

**Figure S3:**
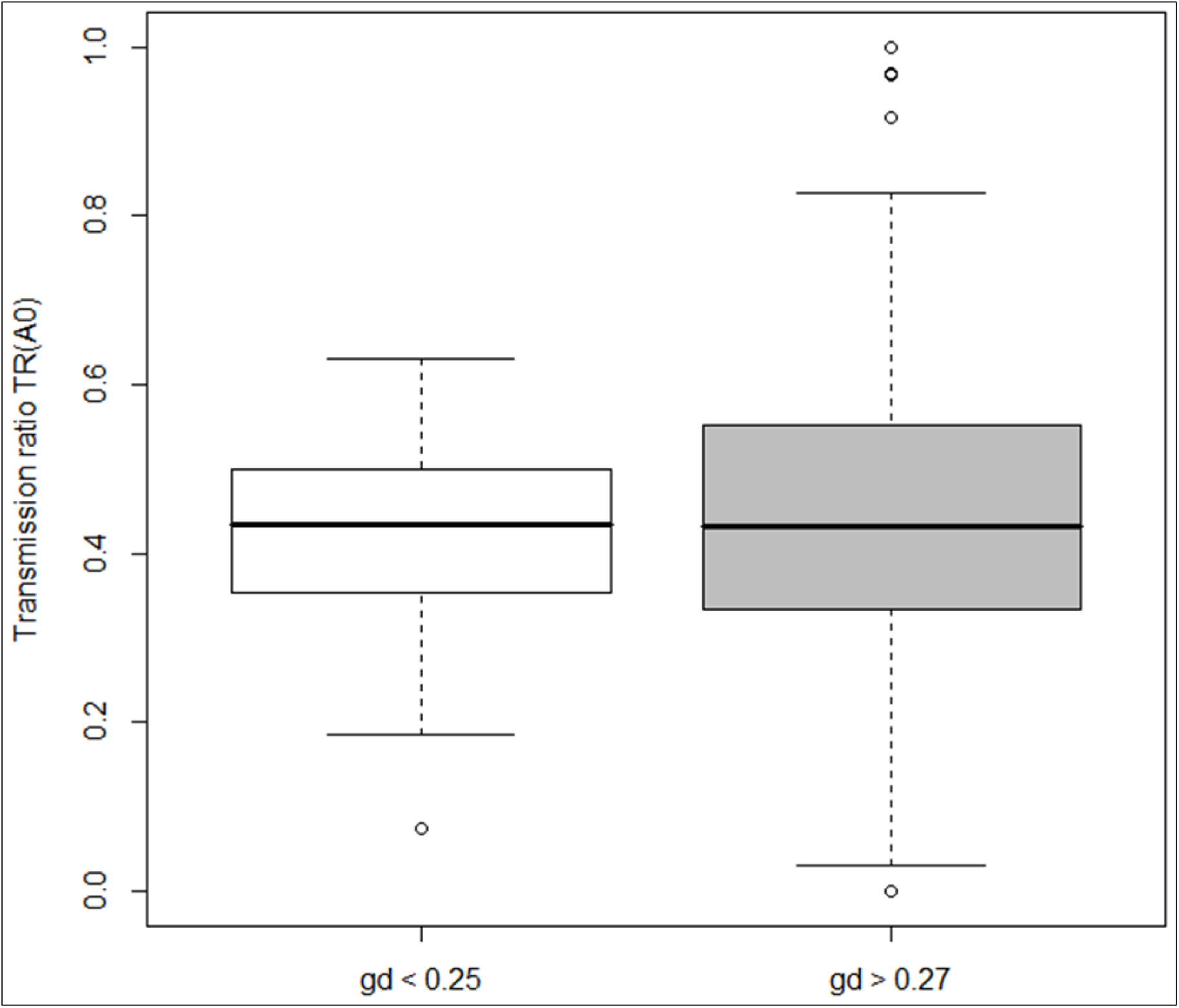
Genetic distance between parents’ effect on copy number transmission ratio distortion (cnTRD) for the F-box gene BT101196. Distribution of transmission ratio values for crosses where the two parents are genetically close (white box) or more distant (grey box).

**Table S1:** Parental and partner genotype effects on copy number transmission ratio distortion (cnTRD).

| GenBank Acc | ♀He x ♂Ho crosses |  |  |  | ♀Ho x ♂He crosses |  |  |  | ♀He x ♂He crosses |  |  |  | He x N crosses |  |  |  | Favored CN | PE | PGE |
| --- | --- | --- | --- | --- | --- | --- | --- | --- | --- | --- | --- | --- | --- | --- | --- | --- | --- | --- | --- |
|  | n | TR(A0) | CI (95%) | P-value | n | TR(A0) | CI (95%) | P-value | n | TR(A0) | CI (95%) | P-value | n | TR(A0) | CI (95%) | P-value |  |  |  |
| BT103424 | 392 | 0.554 | 0.502 - 0.603 | 3.80E-02 | 421 | 0.542 | 0.492 - 0.589 | 9.70E-02 | 308 | 0.601 | 0.560 - 0.639 | 6.60E-07 | 1121 | 0.57 | 0.544 - 0.596 | 1.20E-07 | 0 | D (ME) | I |
| BT109608 | 384 | 0.604 | 0.553 - 0.653 | 5.20E-05 | 387 | 0.543 | 0.491 - 0.593 | 1.00E-01 | 464 | 0.52 | 0.487 - 0.553 | 2.20E-01 | 1235 | 0.544 | 0.520 - 0.568 | 2.70E-04 | 0 | D (ME) | D |
| BT101196 | 407 | 0.536 | 0.485 - 0.584 | 1.70E-01 | 295 | 0.593 | 0.534 - 0.649 | 1.60E-03 | 402 | 0.494 | 0.458 - 0.528 | 7.50E-01 | 1104 | 0.525 | 0.498 - 0.550 | 6.00E-02 | 0 | I (PME) | D |
| BT105201 | 150 | 0.147 | 0.094 - 0.213 | < 2.2e-16 | 213 | 0.249 | 0.192 - 0.312 | 1.10E-13 | NA | NA | NA | NA | 363 | 0.207 | 0.166 - 0.251 | < 2.2e-16 | 1 | I (PME) | ND |
| BT107150 | 276 | 0.236 | 0.186 - 0.290 | < 2.2e-16 | 250 | 0.208 | 0.159 - 0.263 | < 2.2e-16 | 153 | 0.327 | 0.274 - 0.382 | 1.30E-09 | 679 | 0.261 | 0.231 - 0.292 | < 2.2e-16 | 1 | I (PME) | I |
| CO253730 | 150 | 0.353 | 0.277 - 0.435 | 4.10E-04 | 185 | 0.308 | 0.242 - 0.380 | 1.90E-07 | 32 | 0.344 | 0.229 - 0.473 | 1.70E-02 | 367 | 0.331 | 0.284 - 0.379 | 1.30E-11 | 1 | I (PME) | I |
| BT110689 | 319 | 0.386 | 0.331 - 0.441 | 5.20E-05 | 316 | 0.358 | 0.304 - 0.413 | 4.70E-07 | 190 | 0.368 | 0.319 - 0.419 | 3.30E-07 | 825 | 0.37 | 0.340 - 0.400 | < 2.2e-16 | 1 | I (PME) | I |
| BT119696 | 183 | 0.432 | 0.358 - 0.506 | 7.60E-02 | 89 | 0.225 | 0.142 - 0.325 | 1.80E-07 | NA | NA | NA | NA | 272 | 0.364 | 0.306 - 0.424 | 8.50E-06 | 1 | D (PE) | ND |
| BT113426 | 365 | 0.479 | 0.427 - 0.532 | 4.60E-01 | 326 | 0.436 | 0.381 - 0.491 | 2.30E-02 | 31 | 0.468 | 0.339 - 0.598 | 7.00E-01 | 722 | 0.459 | 0.423 - 0.495 | 2.90E-02 | 1 | D (PE) | D |
| BT111788 | 258 | 0.407 | 0.346 - 0.469 | 3.40E-03 | 178 | 0.506 | 0.429 - 0.581 | 9.40E-01 | 249 | 0.446 | 0.401 - 0.490 | 1.70E-02 | 685 | 0.446 | 0.414 - 0.479 | 1.20E-03 | 1 | D (ME) | I |
| BT106719 | 298 | 0.383 | 0.327 - 0.440 | 6.00E-05 | 251 | 0.47 | 0.407 - 0.533 | 3.80E-01 | 86 | 0.424 | 0.349 - 0.501 | 5.60E-02 | 635 | 0.423 | 0.386 - 0.460 | 4.10E-05 | 1 | D (ME) | D |
| BT106118 | 96 | 0.344 | 0.249 - 0.447 | 2.90E-03 | 28 | 0.464 | 0.275 - 0.661 | 8.50E-01 | 30 | 0.4 | 0.275 - 0.534 | 1.60E-01 | 154 | 0.38 | 0.310 - 0.454 | 1.50E-03 | 1 | D (ME) | D |
| DV985927 | 119 | 0.42 | 0.330 - 0.514 | 9.90E-02 | 122 | 0.434 | 0.344 - 0.527 | 1.70E-01 | NA | NA | NA | NA | 241 | 0.427 | 0.364 - 0.492 | 2.80E-02 | 1 | I | ND |
| BT109181 | 87 | 0.414 | 0.309 - 0.524 | 1.30E-01 | 115 | 0.409 | 0.317 - 0.504 | 6.20E-02 | 54 | 0.426 | 0.331 - 0.524 | 1.50E-01 | 256 | 0.416 | 0.360 - 0.473 | 3.70E-03 | 1 | I | I |
| BT116133 | 355 | 0.482 | 0.428 - 0.535 | 5.20E-01 | 244 | 0.484 | 0.419 - 0.548 | 6.50E-01 | 365 | 0.463 | 0.426 - 0.499 | 5.00E-02 | 964 | 0.472 | 0.444 - 0.499 | 4.20E-02 | 1 | I | D |
| DR563872 | 298 | 0.47 | 0.412 - 0.528 | 3.20E-01 | 270 | 0.544 | 0.482 - 0.604 | 1.60E-01 | 94 | 0.367 | 0.298 - 0.440 | 3.30E-04 | 662 | 0.471 | 0.434 - 0.507 | 1.20E-01 | 1 | I | D |
| DR587158 | 29 | 0.414 | 0.235 - 0.610 | 4.60E-01 | 55 | 0.418 | 0.286 - 0.558 | 2.80E-01 | NA | NA | NA | NA | 84 | 0.417 | 0.309 - 0.529 | 1.60E-01 | ND | ND | ND |
| BT103050 | 146 | 0.493 | 0.409 - 0.577 | 9.30E-01 | 146 | 0.432 | 0.349 - 0.515 | 1.20E-01 | 26 | 0.365 | 0.236 - 0.510 | 7.00E-02 | 318 | 0.448 | 0.394 - 0.501 | 5.90E-02 | ND | ND | ND |
| BT119776 | 58 | 0.397 | 0.270 - 0.533 | 1.50E-01 | 119 | 0.504 | 0.411 - 0.597 | 1.00E+00 | NA | NA | NA | NA | 177 | 0.469 | 0.393 - 0.545 | 4.50E-01 | ND | ND | ND |
| BT101202 | 218 | 0.505 | 0.436 - 0.572 | 9.50E-01 | 171 | 0.433 | 0.357 - 0.510 | 9.20E-02 | NA | NA | NA | NA | 389 | 0.473 | 0.422 - 0.523 | 3.10E-01 | ND | ND | ND |
| BT114401 | 272 | 0.467 | 0.406 - 0.528 | 3.00E-01 | 297 | 0.502 | 0.443 - 0.559 | 1.00E+00 | 88 | 0.455 | 0.379 - 0.531 | 2.60E-01 | 657 | 0.478 | 0.441 - 0.514 | 2.40E-01 | ND | ND | ND |
| BT110740 | 84 | 0.476 | 0.366 - 0.588 | 7.40E-01 | 58 | 0.5 | 0.365 - 0.634 | 1.00E+00 | NA | NA | NA | NA | 142 | 0.486 | 0.401 - 0.571 | 8.00E-01 | ND | ND | ND |
| BT109138 | 92 | 0.543 | 0.436 - 0.647 | 4.70E-01 | 61 | 0.443 | 0.315 - 0.575 | 4.40E-01 | 31 | 0.371 | 0.251 - 0.503 | 5.60E-02 | 184 | 0.465 | 0.397 - 0.534 | 3.40E-01 | ND | ND | ND |

He: heterozygote, Ho: homozygote, N: heterozygote or homozygote, CN: copy number allele, PE: parental effect, PGE: partner genotype effect, n: number of offsprings examined, TR(A0): transmission ratio for the copy number allele A0 (zero copy), CI (95%): 95% confidence interval, P-value: *p-value* for two-tailed exact binomial test, NA: not available, ND: not determined, D: dependent, I: independent, (PE): paternal effect, (ME): maternal effect, (PME): paternal and maternal effects.

**Table S2:**
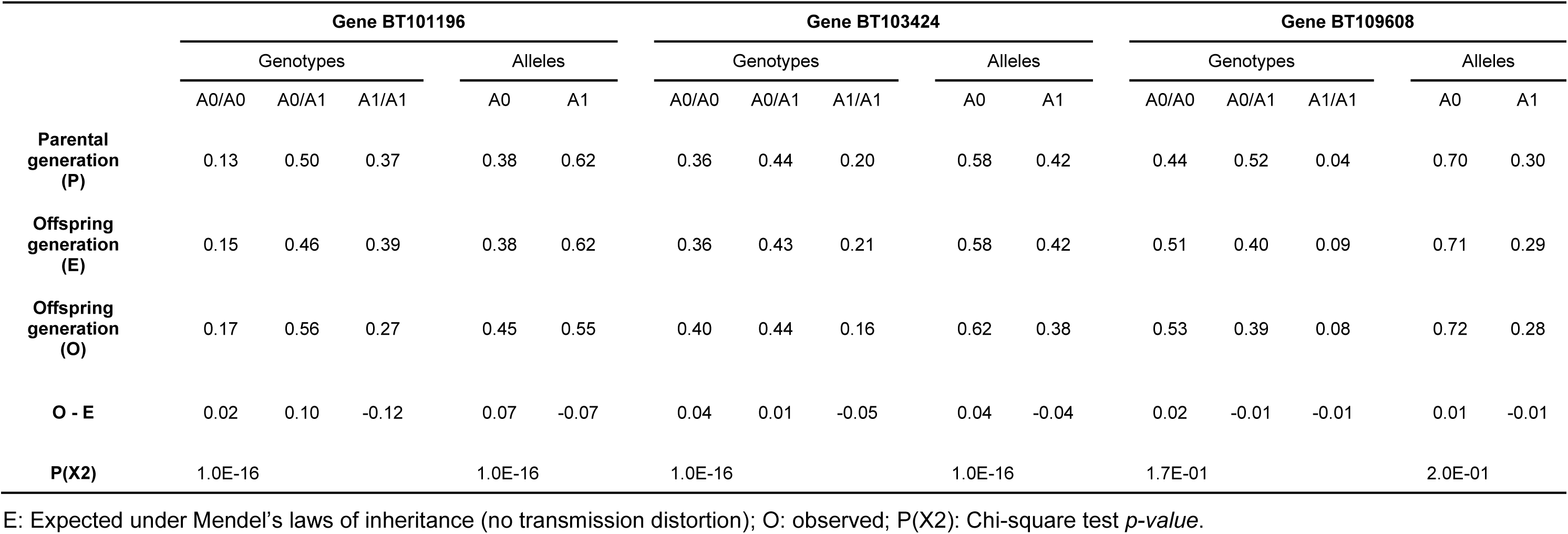
Genotypes and alleles frequencies in the parental and offspring generations for three genes favoring the transmission of zero copy (for all crosses).

|  | Gene BT101196 |  |  |  |  | Gene BT103424 |  |  |  |  | Gene BT109608 |  |  |  |  |
| --- | --- | --- | --- | --- | --- | --- | --- | --- | --- | --- | --- | --- | --- | --- | --- |
|  | Genotypes |  |  | Alleles |  | Genotypes |  |  | Alleles |  | Genotypes |  |  | Alleles |  |
|  | A0/A0 | A0/A1 | A1/A1 | A0 | A1 | A0/A0 | A0/A1 | A1/A1 | A0 | A1 | A0/A0 | A0/A1 | A1/A1 | A0 | A1 |
| Parental generation (P) | 0.13 | 0.50 | 0.37 | 0.38 | 0.62 | 0.36 | 0.44 | 0.20 | 0.58 | 0.42 | 0.44 | 0.52 | 0.04 | 0.70 | 0.30 |
| Offspring generation (E) | 0.15 | 0.46 | 0.39 | 0.38 | 0.62 | 0.36 | 0.43 | 0.21 | 0.58 | 0.42 | 0.51 | 0.40 | 0.09 | 0.71 | 0.29 |
| Offspring generation (O) | 0.17 | 0.56 | 0.27 | 0.45 | 0.55 | 0.40 | 0.44 | 0.16 | 0.62 | 0.38 | 0.53 | 0.39 | 0.08 | 0.72 | 0.28 |
| O - E | 0.02 | 0.10 | -0.12 | 0.07 | -0.07 | 0.04 | 0.01 | -0.05 | 0.04 | -0.04 | 0.02 | -0.01 | -0.01 | 0.01 | -0.01 |
| P(X2) | 1.0E-16 |  |  | 1.0E-16 |  | 1.0E-16 |  |  | 1.0E-16 |  | 1.7E-01 |  |  | 2.0E-01 |  |

E: Expected under Mendel’s laws of inheritance (no transmission distortion); O: observed; P(X2): Chi-square test *p-value*.

**Table S3:**
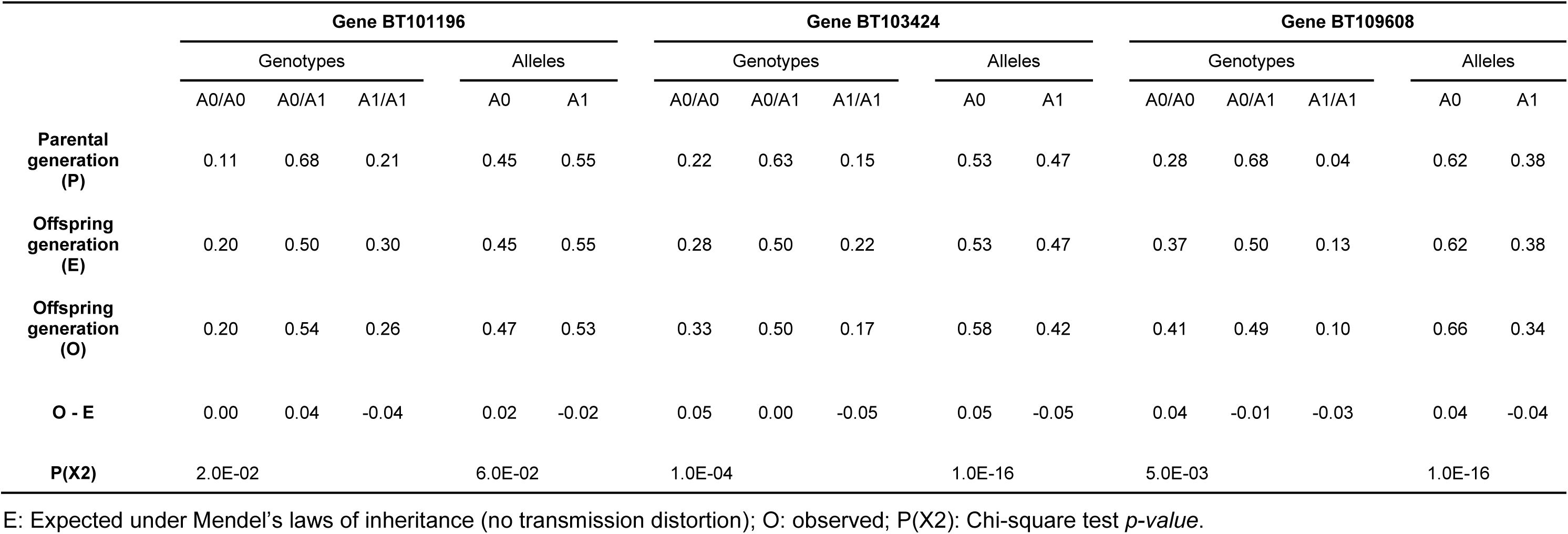
Genotypes and alleles frequencies in the parental and offspring generations for three genes favoring the transmission of zero copy (for crosses with at least one heterozygote parent).

|  | Gene BT101196 |  |  |  |  | Gene BT103424 |  |  |  |  | Gene BT109608 |  |  |  |  |
| --- | --- | --- | --- | --- | --- | --- | --- | --- | --- | --- | --- | --- | --- | --- | --- |
|  | Genotypes |  |  | Alleles |  | Genotypes |  |  | Alleles |  | Genotypes |  |  | Alleles |  |
|  | A0/A0 | A0/A1 | A1/A1 | A0 | A1 | A0/A0 | A0/A1 | A1/A1 | A0 | A1 | A0/A0 | A0/A1 | A1/A1 | A0 | A1 |
| Parental generation (P) | 0.11 | 0.68 | 0.21 | 0.45 | 0.55 | 0.22 | 0.63 | 0.15 | 0.53 | 0.47 | 0.28 | 0.68 | 0.04 | 0.62 | 0.38 |
| Offspring generation (E) | 0.20 | 0.50 | 0.30 | 0.45 | 0.55 | 0.28 | 0.50 | 0.22 | 0.53 | 0.47 | 0.37 | 0.50 | 0.13 | 0.62 | 0.38 |
| Offspring generation (O) | 0.20 | 0.54 | 0.26 | 0.47 | 0.53 | 0.33 | 0.50 | 0.17 | 0.58 | 0.42 | 0.41 | 0.49 | 0.10 | 0.66 | 0.34 |
| O - E | 0.00 | 0.04 | -0.04 | 0.02 | -0.02 | 0.05 | 0.00 | -0.05 | 0.05 | -0.05 | 0.04 | -0.01 | -0.03 | 0.04 | -0.04 |
| P(X2) | 2.0E-02 |  |  | 6.0E-02 |  | 1.0E-04 |  |  | 1.0E-16 |  | 5.0E-03 |  |  | 1.0E-16 |  |

E: Expected under Mendel’s laws of inheritance (no transmission distortion); O: observed; P(X2): Chi-square test *p-value*.

**Table S4:** Genotypes and alleles frequencies in the parental and offspring generations for three genes favoring the transmission of zero copy (for crosses displaying transmission distortions).

|  | Gene BT101196 |  |  |  |  | Gene BT103424 |  |  |  |  | Gene BT109608 |  |  |  |  |
| --- | --- | --- | --- | --- | --- | --- | --- | --- | --- | --- | --- | --- | --- | --- | --- |
|  | Genotypes |  |  | Alleles |  | Genotypes |  |  | Alleles |  | Genotypes |  |  | Alleles |  |
|  | A0/A0 | A0/A1 | A1/A1 | A0 | A1 | A0/A0 | A0/A1 | A1/A1 | A0 | A1 | A0/A0 | A0/A1 | A1/A1 | A0 | A1 |
| Parental generation (P) | 0.00 | 0.50 | 0.50 | 0.25 | 0.75 | 0.17 | 0.72 | 0.11 | 0.53 | 0.47 | 0.42 | 0.50 | 0.08 | 0.67 | 0.33 |
| Offspring generation (E) | 0.00 | 0.50 | 0.50 | 0.25 | 0.75 | 0.28 | 0.50 | 0.22 | 0.53 | 0.47 | 0.43 | 0.50 | 0.07 | 0.68 | 0.32 |
| Offspring generation (O) | 0.00 | 0.60 | 0.40 | 0.30 | 0.70 | 0.34 | 0.51 | 0.15 | 0.59 | 0.41 | 0.51 | 0.45 | 0.04 | 0.74 | 0.26 |
| O - E | 0.00 | 0.10 | -0.10 | 0.05 | -0.05 | 0.06 | 0.01 | -0.07 | 0.06 | -0.06 | 0.08 | -0.05 | -0.03 | 0.06 | -0.06 |
| P(X2) | 1.0E-16 |  |  | 1.0E-16 |  | 1.0E-16 |  |  | 1.0E-16 |  | 7.0E-04 |  |  | 4.0E-04 |  |

E: Expected under Mendel’s laws of inheritance (no transmission distortion); O: observed; P(X2): Chi-square test *p-value*.

